# HuR-Driven Reversible Mitochondrial Shuttling Buffers Cytosolic miRNA Levels in Hepatic Cells to Control Apoptosis

**DOI:** 10.1101/2025.04.08.647748

**Authors:** Saikat Banerjee, Sourav Hom Choudhury, Susanta Chatterjee, Guoku Hu, Kamalika Mukherjee, Suvendra N. Bhattacharyya

**Affiliations:** RNA Biology Research Laboratory, Molecular Genetics Division, CSIR-Indian Institute of Chemical Biology, Kolkata 700032, India; Department of Pharmacology and Experimental Neuroscience, University of Nebraska Medical Center, Omaha, NE 68198-5880, USA; Department of Anesthesiology, University of Nebraska Medical Center, Omaha, NE 68198-4455, USA

**Author notes:** Dipartimento di Biologia, Università degli Studi di Padova, 35121 Padova, Italy (Saikat Banerjee).

**Keywords:** miRNA, mitochondria, HuR, Ago2, miRNA import to mitochondria, mito-miRs

## Abstract

Mitochondria, the “powerhouse” of mammalian cells, also serve as key storage sites for ions, metabolites, and enzymes vital for metabolic regulation. Exploring the regulatory processes that control the activities of miRNAs, the key non-coding RNA in mammalian cells, we found a context-dependent reversible localization of specific miRNAs to the mitochondrial matrix. Our data suggests a *de novo* role of mitochondria as miRNA sinks in mammalian cells. miR-122 is a key hepatic miRNA regulating metabolic processes in the mammalian liver. In this study, we observed increased mitochondrial targeting of miR-122 in amino acid-starved hepatic cells. Interestingly, when cells were refed with amino acids, mitochondrial miR-122 gets relocalized and reused in the cytosol for the translational repression process. Moreover, this phenomenon is not limited to miR-122 as several mitochondrial miRNAs (mito-miRs) follow similar transient storage inside mitochondria in stressed cells. Remarkably, mitochondria-localized mito-miRs preferentially target mRNAs encoding crucial mitochondrial components related to apoptosis. Hence, hepatic cells regulate apoptosis pathways during the starvation-refeeding cycle by shuttling a specific set of miRNAs to and from mitochondria, thereby balancing cytosolic miRNA content and homeostasis. Stress response miRNA binder ELAVL1 or HuR protein was found to be both necessary and sufficient for transporting the mature mito-miRs to the mitochondrial matrix - a process also controlled by the interaction between mitochondria and the endoplasmic reticulum.

**Graphical Abstract:** 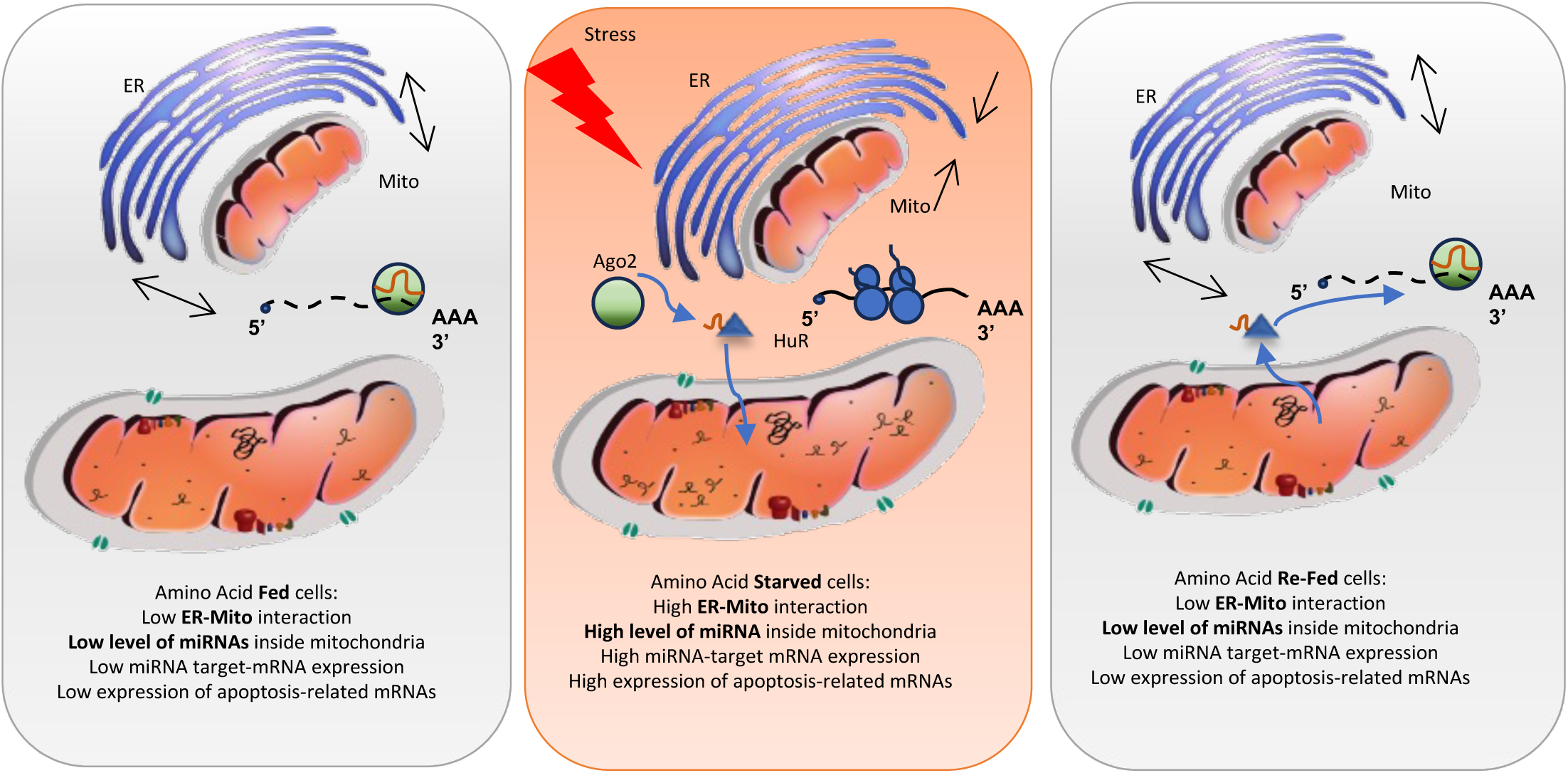

**Key Points:**

- Specific miRNAs (mito-miRs), including miR-122, are targeted to mitochondria for transient storage in stressed hepatic cells.
- Mito-miRs storage in mitochondria allows the expression of apoptosis-related genes in stressed cells.
- Mito-miRs relocalization to cytoplasm on stress reversal allows suppression of apoptosis.
- Binding with ELAVL1 protein HuR allows reversible shuttling of mito-miRs to and from mitochondria in hepatic cells.
- ER-mitochondrial interaction is key for the mitochondrial localization of miRNAs.

## Introduction

miRNAs, the 22-nucleotide-long small RNAs, are critical regulators of cellular processes. miRNAs form complexes with Argonaut proteins to target mRNAs with imperfect binding sites (Bartel, 2009). This interaction reversibly suppresses protein synthesis from specific target mRNAs, a process that is tightly regulated by RNA-binding proteins and miRNA-binding proteins. The activity and abundance levels of miRNA are meticulously regulated in metazoans to ensure effective interactions among mRNA, Ago, and miRNA and proper miRNA function (Pillai *et al*, 2005). Interestingly, exporting miRNAs through extracellular vesicles is essential for sustaining miRNA activity and abundance in metazoan cells (Mukherjee *et al*, 2016). The role of cellular organelles in regulating miRNA activity has recently been observed across diverse cell types (Barman & Bhattacharyya, 2015; Chakrabarty & Bhattacharyya, 2017; Patranabis & Bhattacharyya, 2016; Ray *et al*). The regulation of miRNA binding to Ago proteins represents another mechanism for controlling miRNA activity in mammalian cells (Mazumder *et al*, 2013; Patranabis & Bhattacharyya, 2016). The site-specific phosphorylation of the Ago2 protein is a crucial factor in regulating miRNA activity (Shah *et al*, 2023). This process is essential for uncoupling and inactivating miRNPs, particularly in activated macrophages and differentiating neuronal cells (Mazumder *et al*., 2013; Patranabis & Bhattacharyya, 2016). Furthermore, Ago2-miRNPs can be sequestered in RNA processing bodies (PBs), leading to diminished repressive function due to a lack of interaction with target mRNAs (Mazumder *et al*., 2013; Patranabis & Bhattacharyya, 2016).

Mitochondria, known for their energy production, also regulate miRNA activity (Chakrabarty & Bhattacharyya, 2017; Chatterjee *et al*, 2020). In mammalian cells, a reduction in mitochondrial membrane potential correlates with decreased recruitment of active mTORC1 to the rough endoplasmic reticulum (rER). This observation suggests impaired miRNA recycling and export from cells with reduced mitochondrial potential (Chatterjee *et al*., 2020). Macrophage cells infected with *Leishmania donovani* (*Ld*), a protozoan parasite that causes visceral leishmaniasis, show reduced mitochondrial activity and accumulate higher levels of miRNAs in infected cells (Chakrabarty & Bhattacharyya, 2017). This phenomenon may arise from the inability to effectively target miRNPs to endosomes from the endoplasmic reticulum, hindering the extracellular export and recycling of miRNPs in *Ld*-infected cells.

The growing acknowledgment of mitochondria as reservoirs for diverse metabolites highlights their essential alternative role in cellular function (Nesterov *et al*, 2022). Their interactions with various subcellular structures significantly contribute to membrane biogenesis, apoptosis, and cell cycle checkpoints in diverse mammalian cell types (Audano *et al*, 2020; Gordaliza-Alaguero *et al*, 2019; Xia *et al*, 2019). Metabolic conditions play a significant role in influencing miRNA function and may create a direct link to mitochondrial activity (Carrer *et al*, 2012; Duarte *et al*, 2014; Luo *et al*, 2024). However, the connection between alterations in mitochondrial activity and miRNA function remains largely unexplored.

Researchers have identified specific miRNAs inside mitochondria (Canale & Borghini, 2024; Duroux-Richard *et al*, 2021; Wang *et al*, 2020). However, isolating the “pure” mitochondrial fraction poses a challenge, as the authenticity of mitochondrial purity is complicated by possible contamination from the ER, which hinders the identification of exclusive mitochondria-localized miRNAs (Wang *et al*., 2020). Nevertheless, recent data obtained from “pure” mitochondria with minimal endoplasmic reticulum (ER) contamination suggest a distinct enrichment of specific miRNAs in mammalian mitochondria, some of which are encoded by the mitochondrial genome (Kuthethur *et al*, 2022). Preliminary investigations into a subset of these miRNAs’ associations reveal that mitochondrial localization governs the turnover of specific miRNAs through EV-mediated export (Ma *et al*, 2023a).

Understanding the mechanisms underlying the mitochondrial targeting of nuclear-encoded miRNAs and their transport to the intermembrane space or mitochondrial matrix continues to be a topic of research interest. Additionally, the effects of these mitochondrial-targeted miRNAs on cellular processes are yet to be clarified. Our findings indicate that metabolic conditions modulate the accumulation of mitochondrial miRNAs in a reversible manner. In hepatic cells deprived of amino acids, we observed an accelerated import of miR-122 into the mitochondrial matrix, coinciding with a loss of its interaction with Ago2. Furthermore, we documented increased mitochondrial accumulation of miRNAs in murine liver subjected to fasting or a high-fat diet, suggesting the critical role of mitochondrial miR-122 localization in metabolic regulation processes in mouse liver. To delve deeper into the mechanisms of this localization, we elucidate the significance of ER-mitochondrial contact for accumulating specific miRNAs enriched in mitochondria. Our observations revealed that the ELAVL1 protein HuR, a miRNA-binding protein with a defined role in miRNA derepression and Ago2 uncoupling, is essential for the effective mitochondrial targeting of miRNAs. These findings emphasize the intricate interplay between the intracellular transport of miRNAs to mitochondria and cellular metabolism processes.

Notably, miRNAs from mitochondria relocalize upon refeeding amino acids in hepatic cells, resulting in miRNA reloading with Ago2 in the cytoplasm to repress their respective target genes. Intriguingly, mitochondria-localized miRNAs (mito-miRs) have preferential target sites in mRNAs transcribed from the nuclear genome, which encodes various mitochondrial proteins involved in apoptosis. This indicates a regulatory role for mitochondrial miRNAs in controlling apoptosis in mammalian cells during the starvation-refeeding cycle. Thus, mitochondria can act as miRNA sinks in mammalian cells.

## Results

### Specific miRNAs are localized to the mitochondrial matrix of mammalian cells

The presence of miRNA in the crude or impure mitochondrial fraction must be confirmed in pure mitochondria isolated from mammalian cells free from substantial ER and cytoplasmic contamination (Liao *et al*, 2020) to conclude on exclusive localization of specific miRNAs inside mitochondria. To isolate pure mitochondria, HeLa cells were lysed in an isotonic lysis buffer that was followed by differential multistep separation of organelles to get “pure” mitochondrial fraction free from cytosolic and other organellar contaminants (**Fig. 1A**). In the “pure” mitochondrial fraction, we observed enrichment of mitochondrial protein Cytochrome C and have detected other mitochondrial proteins like VDAC and FACL4. The isolated pure mitochondria are free from calnexin and β-tubulin-two markers for ER and cytoplasmic fractions, respectively (**Fig.1 B**). We have also isolated the mitochondria-associated membrane (MAM) comprising both ER and mitochondria structures, which was positive for mitochondrial markers and ER marker protein calnexin. The MAM fraction is free from cytoplasmic contamination as β-tubulin was not detected there (**Fig.1B**).

**Figure 1.**
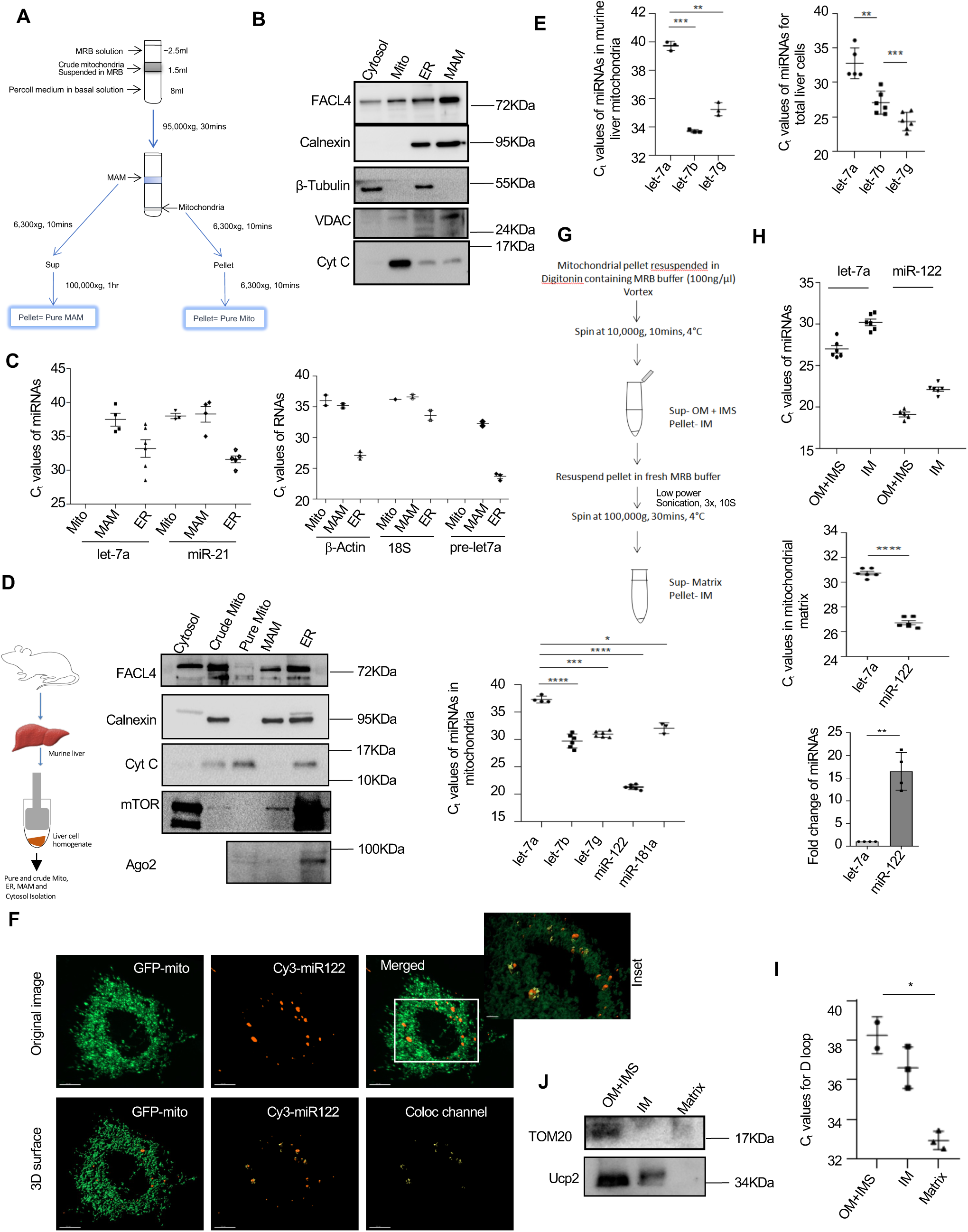
Mitochondrial targeting of miRNAs in mammalian cells. **A.** Schematic representation of isolating pure mitochondria (pure mito) and mitochondria-associated membranes (MAMs) from the crude mitochondrial lysate of HeLa cells via differential and gradient ultracentrifugation. The technique outlines the relative amounts of lysate and solution, alongside details of organelle localization tendencies and centrifugation speed and time during the process isolation. **B.** Western blots illustrate the characterization of fractions obtained after subcellular isolation, with protein blots indicating specific organelle markers in HeLa cells. The positions of the molecular weight markers are indicated along the side of each panel. **C.** qRT–PCR of miRNAs and non-miRNAs was performed using RNA isolated from subcellular fractions (pure mitochondria, ER, and MAM). Relative Ct values indicate the abundance of 18S, pre-let-7a, and β-actin in different fractions. Ct values also show the relative abundance of let-7a and miR-21 in various subcellular compartments. All measurements were taken with 200 ng of RNA (n = 3). **D.** The schematic illustrates the isolation of mitochondria, membrane-associated mitochondria (MAM), and pure mitochondria from murine liver. Western blots display mitochondrial and non-mitochondrial proteins in isolated subcellular fractions. The right panel shows the Ct values for multiple miRNAs in RNA isolated from the pure mitochondrial fraction. The lowest Ct value was detected for miR-122, indicating its enrichment in liver-isolated mitochondria (n =3) **E.** qRT–PCR data reveal the relative abundance, denoted by Ct values, of the let-7 miRNA family members let-7a, let-7b, and let-7g in total liver lysate and liver-derived mitochondria-associated RNAs. For total RNA estimation, 200 ng of total cellular RNA and 30 ng of mitochondrial RNA were used for PCR analysis (n = 3). **F.** Huh7 hepatic cells transfected with Cy3-3’ tagged miR-122 were imaged to observe its distribution within the transfected hepatic cells. The captured confocal image (60X) and 3D surface rendering of the channels are presented, with yellow indicating areas of colocalization. The inset displays a 2X magnified view to emphasize colocalization. Bars measure 10µm. **G-H.** A schematic representation of the separation strategy used for mitochondrial inner and outer membranes and matrix fractions involves detergent solubilization, centrifugation, and sonication (G). Ct value analysis via qRT-PCR for miR-21 and let-7a in purified sub-mitochondrial fractions (n> 3) confirms the presence of miRNAs within the mitochondrial matrix. Data is normalized against Ct values obtained from the mitochondrial D-loop (n> 3). A several-fold enrichment of miR-122 was detected compared to let-7a in the mitochondrial matrix (H). **I-J.** Mitochondrial D-loop RNA estimated through relative Ct analysis demonstrates the effectiveness of the isolated mitochondrial matrix fraction as illustrated in panel F (n=3) (I). Western blots for TOM 20 confirm its presence in the outer membrane, while Ucp2 is found in both membranes (J). Data are expressed as mean ± SD; ns denotes nonsignificant, ∗p < 05, ∗∗p < 01, ∗∗∗p < 001, ∗∗∗∗p < 0001, based on a two-tailed Student’s t-test. MRB stands for Mitochondrial Resolution Buffer; MAM refers to Mitochondria-associated membrane.

Mitochondria has its own translational machinery to express the mitochondrial genome-encoded proteins inside the mitochondrial matrix (Wang *et al*, 2021). Still, the association of nuclear-encoded mRNAs and 80S ribosomes with the outer surface of the mitochondrial outer membrane (OM) has been reported before (Sylvestre *et al*, 2003). Consistent with this notion, we detected the presence of 18S ribosomal RNA and β-Actin mRNA with purified mitochondria from HeLa cells (**Fig. 1C**). However, neither pre nor mature let-7a was detected in the RNA isolated from the mitochondrial fraction, although both were present in the ER and MAM fractions. In contrast to let-7a, miR-21 was detected in the pure mitochondria and MAM fractions. Both let-7a and miR-21 were abundantly present in the ER fraction (**Fig.1C**). We aimed to determine whether we could detect miRNAs in mitochondria purified from mammalian tissues. We used mouse liver lysate to purify mitochondria and verified the purity by performing western blot analysis for Cytochrome C and FACL4 as mitochondrial markers. We found no ER or cytoplasmic contamination in the purified mitochondria, as we did not detect calnexin or mTOR in the pure mitochondrial fraction. Ago2 is also largely absent from the pure mitochondrial fraction (**Fig.1D**). Comparative Ct value analysis suggests an abundance of the liver-specific miR-122 in liver-derived mitochondria (>106 molecules/ng of mitochondrial RNA isolated) (**Fig.1D**). Among the three members of the let-7 family of miRNAs that were checked for their abundance in murine mitochondria, let-7a was found to be minimal, while let-7b is relatively more enriched in mouse liver mitochondria (**Fig.1E**). miR-122 is the most abundant miRNA in the liver and regulates many metabolic pathways in liver cells (Wen & Friedman, 2012). To confirm the mitochondrial association of miR-122, we have done an immune fluorescence analysis of Mito-GFP-expressing cells transfected with Cy3-tagged miR-122. The mitochondria are tagged with GFP, and we detected colocalization of the Cy3-miR-122 with GFP-positive mitochondria (**Fig. 1F**).

It is essential to determine whether the miRNA detected in the mitochondrial fraction is solely associated with the outer mitochondrial membrane or directed to the mitochondrial matrix. To address this question, we conducted a mitochondrial sub-fractionation experiment to isolate the outer mitochondrial membrane using digitonin treatment, resulting in mitoplasts containing only the inner membranes and matrix, excluding the outer membrane and intermembrane space content (Sileikyte *et al*, 2011). We have subsequently lysed mitoplast by sonication to separate the insoluble inner membrane from soluble matrix components (Huang *et al*, 2024). Analyzing the RNA isolated from equal amounts of outer membrane and intermembrane space contents, we found the presence of miRNAs there. However, a substantial amount of miRNA was identified as being targeted to the inner membrane. By comparing other mitochondrial compartments, we detected that miR-122 was abundant in the mitochondrial matrix (Fig. **1G and H**). The effectiveness of mitochondrial fractionation in separating the outer membrane (along with the intermembrane space fraction), inner membrane, and mitochondrial matrix was confirmed by the presence of the mitochondrial outer membrane marker protein TOM20 and inner membrane-localized Ucp2 proteins in differentiated sub-mitochondrial fractions. TOM20 and Ucp2 proteins were absent in the matrix fraction, while the amplification of the D loop region from mitochondrial DNA confirms its exclusive presence in the mitochondrial matrix fraction (**Fig. 1 I and J).**

### Mitochondrial-ER interaction is essential for the mitochondrial targeting of miRNAs

ER membranes are the sites where miRNA-mediated translation repression of mRNAs occurs before the repressed mRNAs are targeted to endosome fractions for subsequent degradation of target mRNAs and recycling of miRNAs detached from target mRNAs (Bose *et al*, 2020). The mitochondria-associated membrane (MAM) is a specialized ER structure attached to mitochondria (Liu & Yang, 2022; Missiroli *et al*, 2018), and we have noted the association of MAM with miRNAs but not with Ago2 (**Fig.1 C and D**). Mfn1 and Mfn2 facilitate the interaction between mitochondria and the endoplasmic reticulum (ER). Among these, Mfn2 plays a significant role in the mitochondria-ER interaction, and impairment of Mfn2 has been reported to be associated with an enhanced accumulation of miRNA within the ER fraction (Bose *et al*., 2020; Chakrabarty & Bhattacharyya, 2017; Chatterjee *et al*., 2020; Wang *et al*., 2020). We hypothesize that miRNAs uncoupled from Ago2 associate with MAM before being transferred to the mitochondrial matrix. Thus, we aimed to examine the effect of Mfn2 depletion on the mitochondrial association of miRNAs. Using mouse embryonic fibroblast (MEF) cells from both wild-type and Mfn2-/- mice, characterized by a documented loss of mitochondria-ER interaction (Chakrabarty & Bhattacharyya, 2017), we observed a diffused distribution of mitochondria in the cytoplasm (**Fig. 2A and B**). With isolated mitochondria and MAM from wild-type and Mfn2-/- cells, we have observed a change in the association of FACL4, the mitochondrial marker protein, with MAM, indicating a loss of ER-mitochondria interaction (**Fig. 2C**). The Mfn2 depletion is associated with enhanced cellular miRNA levels (Chakrabarty & Bhattacharyya, 2017). However, a decrease in miRNA content in mitochondria is observed in most miRNAs in Mfn2 KO MEF isolated pure mitochondria (**Fig. 2D)**. Interestingly, when we downregulated Mfn2 by siRNAs in Huh7 cells with siRNAs to reduce Mfn2 expression, we also noted a decrease in mitochondrial targeting of miRNAs (**Fig. 2 E and F**).

**Figure 2.**
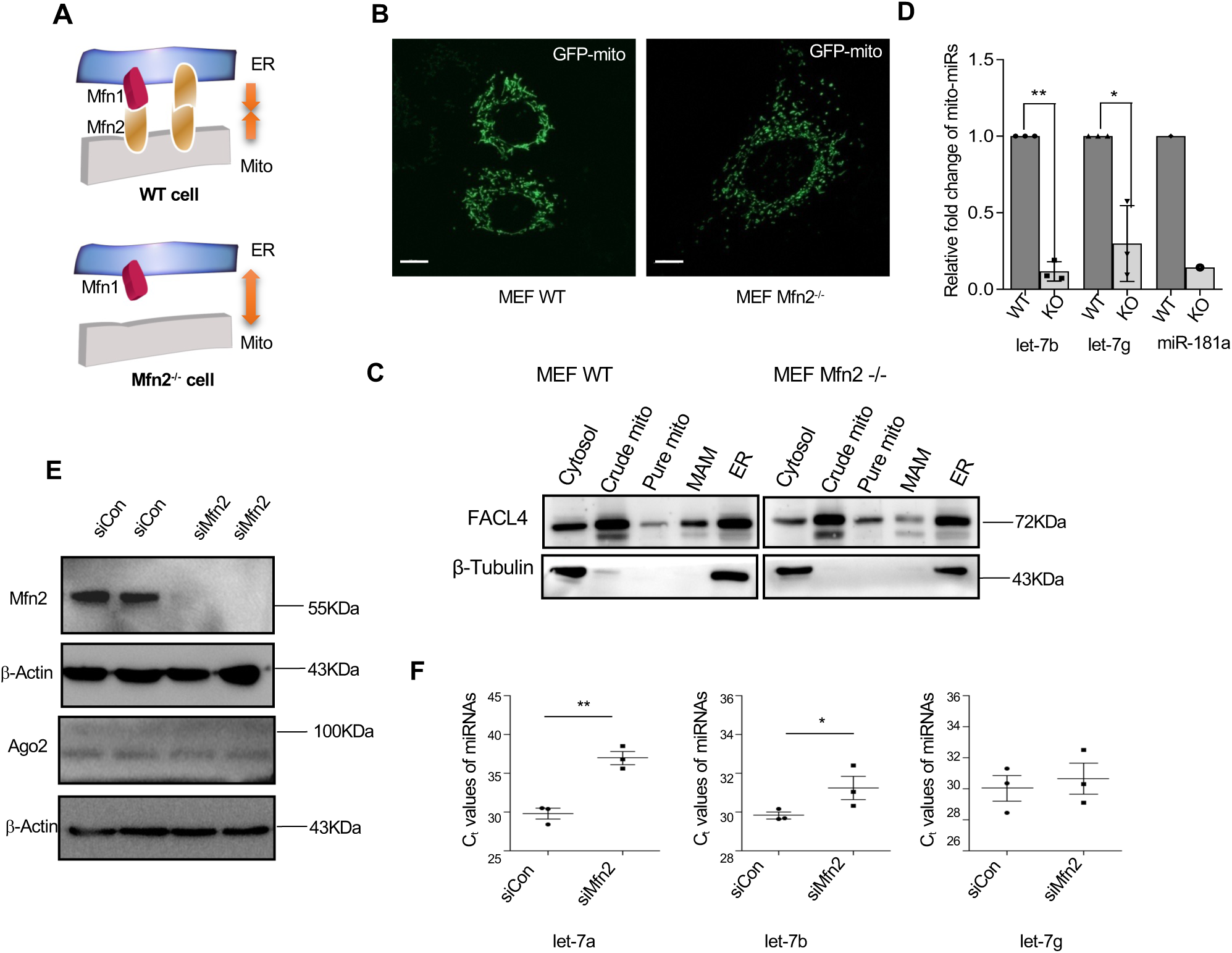
Mitochondria ER-Interaction is required for miRNA targeting to mitochondria. A-B. Cartoon representation of mitochondrial membrane tethering with endoplasmic reticulum (ER) membranes mediated by mitofusin-2 (Mfn2). In wild-type cells, the ER and mitochondria are connected through Mfn2, forming homo- or hetero-dimers with Mfn-1 to maintain their contact. These connections are lost under Mfn2-/- knockout conditions, leaving the ER and mitochondrial membranes separate (A). Confocal images show GFP-tagged mitochondrial structures in wild-type and Mfn2-/- MEF cells (B). Bars measure 10µm. **C.** Western blot analysis of FACL4 and β-tubulin in mitochondria, MAM, and other fractions isolated from wild-type (WT) and Mfn2-/- MEF cell lysates confirms the reduction of FACL4 associated with Mfn2-/- isolated MAM. **D.** Comparative Ct values of miRNAs in pure mitochondrial fractions isolated from WT and Mfn2 Knockout (KO) cells (n> 3) indicate a loss of the let-7 family of miRNAs that are otherwise enriched in mitochondria. The fold decrease of let-7b and let-7g mito-miR abundance in Mfn2 KO cells is shown in the lower panel, normalized against the let-7a level **E.** Western blot data indicate siRNA-mediated silencing of Mfn2 in hepatic Huh7 cells. The lower panels display Ago2 levels in Mfn2 knockdown cells, with β-Actin serving as the loading control **F.** Ct values of miRNAs from pure mitochondrial fractions (n> 3) show a loss of the let-7a family of miRNAs in Mfn2-silenced Huh7 mitochondria, with higher Ct values for let-7a, let-7b, and let-7g compared to control siRNA-treated mitochondria Data are presented as mean ± SD; ns indicates nonsignificant; p < 0.05, p < 0.01, p < 0.001, and p < 0.0001 were calculated using a two-tailed Student’s t-test.

### Enhanced mitochondria-ER interaction augments mitochondrial targeting of miRNAs in amino acid-starved cells

Amino acid starvation is a mechanism that decreases the cellular levels of specific miRNAs by promoting their export through extracellular vesicles to ensure the expression of mRNAs targeted by miRNAs that are essential for the stress response (Mukherjee *et al*., 2016). miR-122 is the predominant miRNA in hepatic cells that showed enhanced export in amino acid-starved hepatocytes (Mukherjee *et al*., 2016). The increase in miRNA export does not explain the non-functional nature of the residual miRNAs left in stressed hepatic cells with reduced translational repressive activity (Mukherjee *et al*., 2016). There must be multiple mechanisms to regulate miRNA function that define the inactive miR-122 present in amino acid-starved hepatic cells. We have observed an increased number of RNA processing bodies in Huh7 cells that were starved of amino acids for 16 hours, confirming a change in subcellular structures, which are known to be altered during starvation, where miRNA-repressed mRNAs are also localized in control amino acid-fed cells (Bhattacharyya *et al*, 2006) (**Fig.S1A-C**). Amino acid starvation also enhanced the mitochondrial volume and surface area (**Fig.S1 D-F**). Using 3D structure reconstruction and analysis, we documented an enhanced mitochondrial tubular network in starved cells (**Fig.S1 E**). We observed increased interaction between mitochondria and the endoplasmic reticulum (ER) in amino acid-starved Huh7 cells, along with a moderate rise in Mfn2 expression levels (**Fig. S1G-I)**. Interestingly, although Ago2 is not associated with pure mitochondria in HeLa cells (**Fig. 1D**), we noted the proximal localization of Ago2-positive bodies with mitochondria during microscopic analysis, which increases in starved Huh7 cells (**Fig. S2**) We have noted that loss of mitochondria-ER interaction due to Mfn2 depletion caused defective miRNA accumulation in mitochondria (**Fig.2D and F**). How does the enhancement of mitochondrial-ER interaction in starved Huh7 cells affect the mitochondrial miRNA content? We measured the miRNA content of mitochondria isolated from fed and amino acid-starved Huh7 cells to document the increased levels of mito-miRs in mitochondria from starved hepatic cells. In contrast, the overall cellular miRNA content was reduced (Mukherjee *et al*., 2016) (**Fig. 3 A-B**). The altered expression of phosphorylated eIF2α and phospho-eIF4E-BP1 confirmed the induction of stress in Huh7 cells that were starved of amino acids for 16 hours (**Fig.3 C**). How is the mitochondrial targeting of mito-MiRs affected in the *in vivo* setting? We starved C57BL/6 mice of amino acids by placing them on a glucose-only diet for 12 hours (Mukherjee *et al*., 2016) to observe a similar decrease in the miRNA content of hepatocytes alongside an increased level of mitochondrial miRNA content in the livers of starved mice (**Fig. 3 D-E**). Exploring the subcellular localization of specific miRNAs, we found that mito-miR, although present in both sub-mitochondrial fractions, let-7g is specifically localized more with the inner membrane and matrix-associated fraction (**Fig 3F-G**).

**Figure 3.**
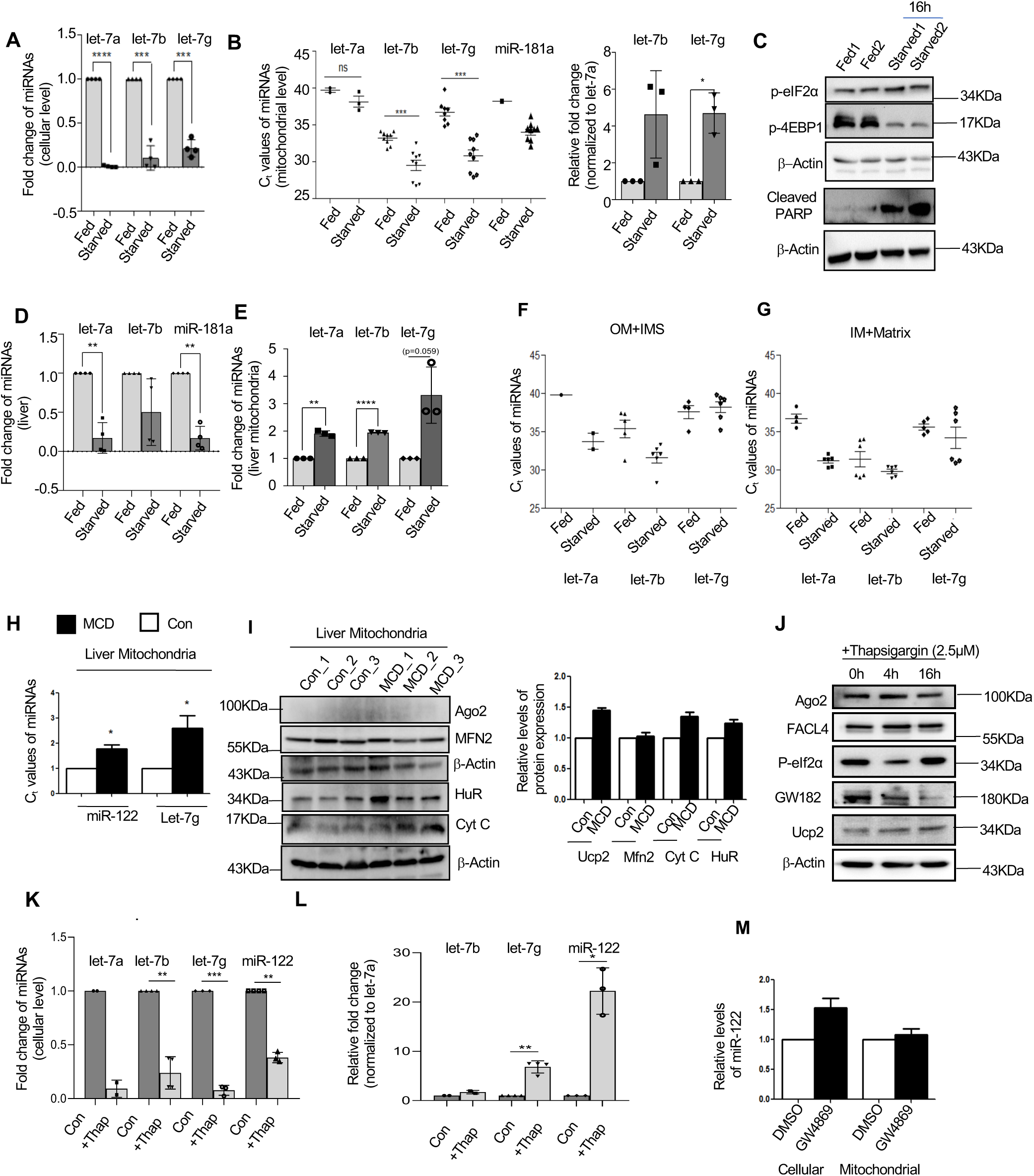
The Starvation-induced enrichment of mito-miRs in hepatic cell mitochondria. **A.** Fold change in cellular miRNA content after 16 hours of amino acid starvation of Huh7 cells was analyzed. Relative levels of various miRNAs were quantified using qRT–PCR, and the data were normalized to U6 snRNA (n> 3). **B.** miRNA-specific Ct values are plotted to illustrate the changes in their relative abundance in purified mitochondria following amino acid starvation of Huh7 cells. The analysis shows a significant decrease in Ct values for the miRNAs let-7b, let-7g, and miR-181a in the mitochondria of starved cells, indicating their enrichment in the mitochondrial fraction during starvation. A bar graph presents the fold enrichment of let-7b and let-7g mito-miRs in purified mitochondria compared to let-7a **C.** Western blot analysis performed on fed and starved Huh7 extracts showed increased levels of the stress marker protein phospho-eIF2α and cleaved PARP, along with decreased phosphorylation of eIF-4EBP1. β-actin served as the loading control for all samples. **D.** Cellular miRNA levels in the liver of mice vary under a normal diet (Fed) and after 12 hours of starvation, as estimated by qRT–PCR and normalized against U6 snRNA levels (n=3). **E.** The fold changes of let-7b and let-7g in the isolated pure mitochondrial fraction obtained from the livers of mice on a normal diet (Fed) or starved for 12 hours. Levels of each miRNA in the Fed condition are considered as units. **F-G.** The relative enrichment of mito-microRNAs in the outer and intermembrane space (F) as well as in the inner membrane and mitochondrial matrix (G) of mitochondria isolated from control (fed) and 16-hour amino acid-starved Huh7 cells. Relative Ct values are plotted. **H.** Folds change in mitochondrial miR-122 and let-7g levels in the liver mitochondria of animals exposed to a control and Methionine Choline Deficient (MCD) diet (n=4). **I.** Western blot analysis (left panel) and relative quantification (right panel) of different proteins in the mitochondrial fractions isolated from control and MCD diet-exposed animal livers **J.** The effect of Thapsigargin on the expression of stress-responsive proteins in Huh7 cells was determined by western blot analysis, using β-actin as a loading control. **K-L.** The effect of Thapsigargin treatment on cellular (K) and mitochondrial (L) miRNA content was analyzed. Relative levels of miRNAs were measured and plotted. U6 and let-7a were used for normalization in both cellular and mitochondrial samples. **M**. Cellular miR-122 levels increased, while mitochondrial levels remained unchanged when cells were treated with a pharmacological blocker of extracellular vesicle production GW4869. Normalization was performed using U6 snRNA and let-7a for cellular and mitochondrial miR-122, respectively. Statistically analyzed data is presented as mean ± SD from three experiments, with significance levels indicated as ns (nonsignificant), ∗p < 0.05, ∗∗p < 0.01, ∗∗∗p < 0.001, and p < 0.0001, calculated using the two-tailed Student’s t-test.

The methionine choline-deficient (MCD) diet-induced model of NASH is a well-established model in which miRNA dysregulation has been reported in the hepatic cells of mice exposed to the MCD diet (Bandyopadhyay *et al*, 2023; Bandyopadhyay *et al*, 2020). With a high metabolic burden, we also observed enhanced localization of mito-miRs in the mitochondria of liver cells isolated from MCD diet-fed animals (**Fig.3H-I**).

Does enhanced mito-MiR entry into mitochondria occur due to an improved stress response in starved Huh7 cells? Thapsigargin (TG) induces stress that affects cellular miRNA content. We treated Huh7 cells with TG to induce ER stress (Mukherjee *et al*., 2016). TG is also recognized for enhancing the interaction between mitochondria and the endoplasmic reticulum (ER) (Bravo *et al*, 2011). Upon TG treatment, we observed a significant decrease in cellular miRNA content in Huh7 cells, while Ucp2 levels increased. In contrast, mitochondrial miRNA levels were elevated following TG treatment, and western blots confirmed TG’s impact on proteins associated with this pathway (**Fig 3J-L**).

How is mitochondrial miRNA storage in stressed cells linked to its extracellular export? Interestingly, treating cells with GW4869, the blocker of EV-mediated miRNA export from animal cells (Ghosh *et al*, 2015; Mukherjee *et al*., 2016), failed to show any increase in mitochondrial miRNA content. In contrast, total cellular miRNA content was enhanced (**Fig. 3M**). This indicates a lack of crosstalk between the EV-mediated miRNA export pathway and the mitochondrial targeting of mito-miRs.

### miRNA-binding protein HuR is necessary and sufficient for miRNA-targeting to mitochondria

We have seen such a decrease in mitochondrial miRNA content in cells treated with H_2_O_2_ -an inducer of oxidative load on mitochondria and a known agent to increase mitophagy (Zhang *et al*, 2021). We documented enhanced interaction between mitochondria and lysozymes, as well as a large number of lysosomes in H2O2-treated cells. To rule out the possibility that defective mitophagy in starved hepatic cells accounts for the excess accumulation of miRNA-loaded mitochondria, we conducted experiments to evaluate the mitophagy status of the starved Huh7 cells. An increased number of lysosomes was observed in amino acid-starved cells, suggesting a positive effect on mitophagy during starvation (**Fig.S3**). Therefore, enhanced mito-miRs inside the mitochondria of starved cells are not due to defective mitophagy in stressed hepatic cells.

In MCD diet mice liver, we have noted increased expression of the protein HuR (**Fig. 3I**). HuR is a stress-responsive protein that is both necessary and sufficient for miRNA activity derepression and extracellular export process in hepatic cells observed under amino acid starvation (Bandyopadhyay *et al*., 2023; Bhattacharyya *et al*., 2006; Ghosh *et al*, 2024; Goswami *et al*, 2020; Kundu *et al*, 2012). HA-HuR expression can induce the derepression of miRNA-targeted messages having HuR binding sites (Bhattacharyya *et al*., 2006). HuR uncouples miR-122 from Ago2 and facilitates its export from starved hepatic cells, a step crucial for optimal stress response in these cells (Bhattacharyya *et al*., 2006; Mukherjee *et al*., 2016). Interestingly, under stress, the residual cellular miR-122 should remain in a non-functional condition to allow the expression of its target genes in hepatic cells (Bhattacharyya *et al*., 2006). We have documented the accumulation of Ago2-uncoupled miRNAs in the mitochondria of starved cells. Does HuR-mediated miRNA-Ago2 uncoupling also lead to targeting HuR-bound miRNAs to the mitochondria? With HA-HuR expression, we found an enhanced interaction of ER with mitochondria (**Fig. 4A-B**), while HuR depletion resulted in fragmented mitochondrial structure as observed in Mfn2 negative cells (**Fig.4C**). We have observed an increase in HuR expression after 16 hours of starvation in Huh7 cells. Additionally, we found a substantial amount of HA-HuR in the mitochondrial fraction, suggesting a potential role for HuR in mitochondrial function (**Fig. 4 D-E**). The isolation and quantification of mitochondria-associated miRNAs from control and HA-HuR expressing cells indicate enhanced entrapment of miRNA in mitochondria upon ectopic HA-HuR expression. Conversely, the depletion of HuR by siRNA has the opposite effect on the mitochondrial content of miRNAs in hepatic Huh7 cells (**Fig. 4F**)

**Figure 4.**
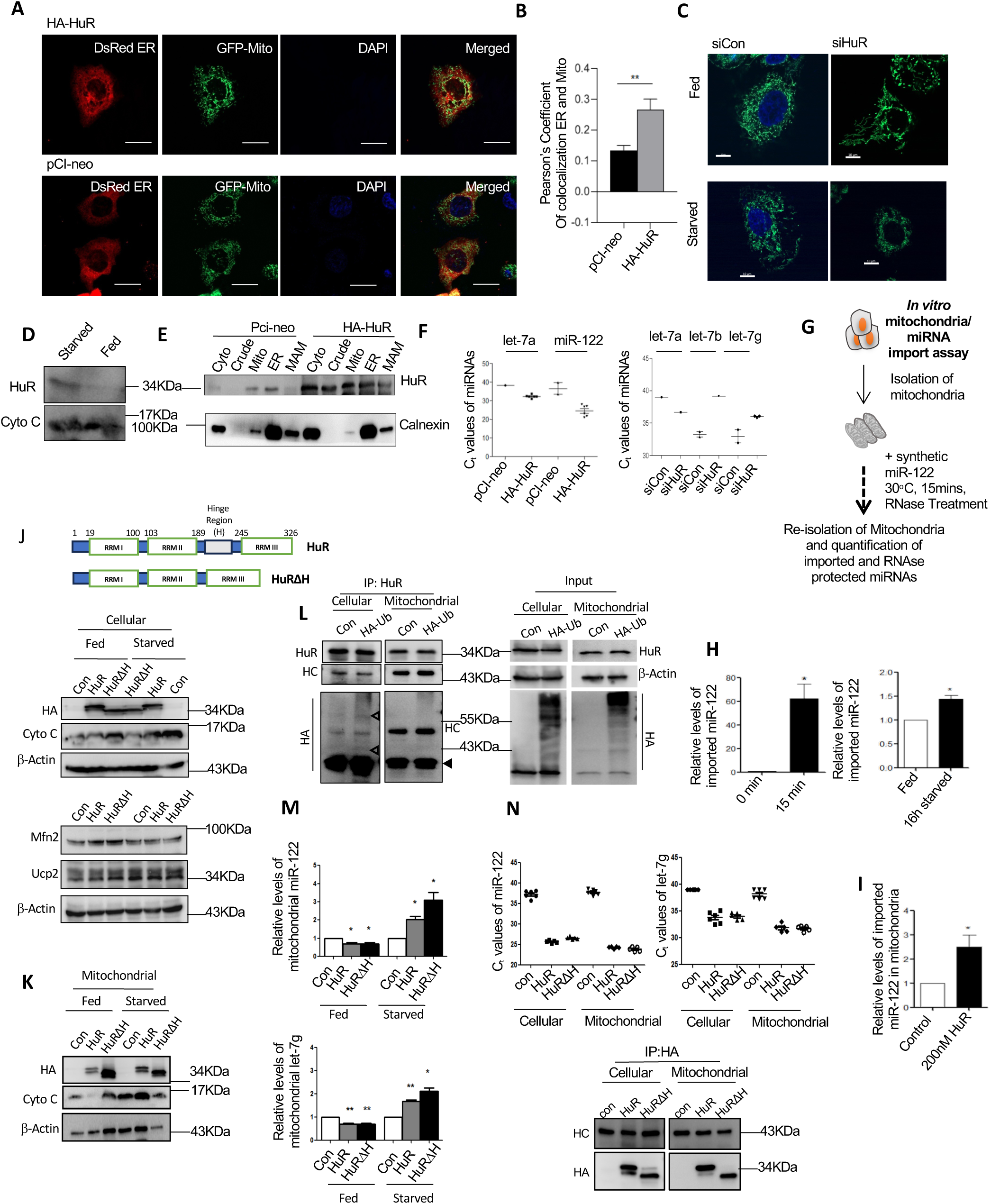
HuR is necessary and sufficient for the mitochondrial targeting of mito-miRs. **A.** Confocal images of Huh7 cells expressing GFP-Mito and DsRed-ER were transfected with either the pCI-neo or HA-HuR expression plasmid and starved for amino acids for 16 hours. **B.** Pearson’s colocalization coefficient for DsRed ER and GFP-Mito in Huh7 cells (n=3), transfected with either pCI-neo or HA-HuR expression plasmids, and starved for 16 hours hrs. **C.** Changes in mitochondrial morphology in GFP-Mito-expressing Huh7 cells transfected with siRNAs specific to HuR (siHuR) or control siRNAs (siCon) under fed conditions and 16 hours of amino acid starvation conditions. **D.** The impact of 16 hours of amino acid starvation on cellular levels of HuR and Cytochrome C was assessed through western blot analysis. **E.** Subcellular distribution of HuR in cells transfected with either pCI-neo or HA-HuR expression plasmids. Subcellular fractions were separated using differential centrifugation and density gradient methods as described in Figure 1. Individual fractions were then processed for protein precipitation and western blotting for HuR and Calnexin (ER) marker). **F.** The difference in Ct values from qRT-PCR analysis of RNA isolated from the mitochondria of Huh7 cells transfected with either the pCIneo or HA-HuR expression plasmid (left panel), or from cells transfected with control and HuR-specific siRNAs (right panel) (n=3). **G.** A flowchart of the in vitro assay for mitochondrial targeting of miRNA was created using isolated mitochondria from Huh7 cells. The mitochondria were incubated with single-stranded miR-122 as substrates, followed by the degradation of non-imported miRNAs inside the mitochondrial matrix. The RNase-protected miRNAs are recovered and quantified. **H.** Following the in vitro reaction and RNase protection, the RNases protected miR-122 was recovered and quantified using qRT-PCR. The related changes in the amount of protected RNA were plotted against the incubation time (left panel) and starvation (right panel). Mitochondrial let-7b levels were utilized for normalization. **I.** Effect of 200 nM recombinant HuR addition on the mitochondrial import of miR-122 in vitro. Internal let-7b was utilized for normalization. **J.** The upper panel presents a diagrammatic view of the full-length and truncated HuR (which is defective for ubiquitination due to the deleted hinge region). The lower panels show expression levels of mitochondrial and non-mitochondrial proteins, as determined by western blots of cellular extracts from Huh7 cells expressing the pCI-neo control, HA-HuR, or the HuR deletion mutant HA-HuRι1H, under fed conditions and after 16 hours of amino acid starvation. **K.** Protein levels were determined by western blots of pure mitochondrial extracts isolated from fed and starved Huh7 cells expressing pCI-neo control, HA-HuR, or HuR deletion mutant HA-HuRι1H. **L.** The presence of ubiquitinated HuR, indicated by hollow arrowheads, was identified in cellular extracts and is absent in mitochondria-associated fractions. This was determined using HA-specific antibodies in immunoprecipitated materials obtained with HuR-specific antibodies, or in inputs from extracts used for immunoprecipitation. A filled arrowhead denotes non-ubiquitinated HuR. HC: Heavy Chain. **M.** Relative changes in mitochondrial miRNA content in fed and starved Huh7 cells expressing pCI-neo-Control, HA-HuR, and HA-HuR ΔH expression plasmids. **N.** The levels of miRNAs associated with HA-immunoprecipitated materials from mitochondrial and cellular extracts of starved Huh7 cells expressing pCI-neo Control, HA-HuR, and HA-HuR ΔH expression plasmids were analyzed. Confirmation of the immunoprecipitation of HuR and its truncated version is shown at the bottom panel. Statistically analyzed data is presented as mean ± SD, with significance indicators: ns (nonsignificant), ∗p < 0.05, ∗∗p < 0.01, ∗∗∗p < 0.001, and p < 0.0001, calculated using a two-tailed Student’s t-test.

How does HA-HuR facilitate the entry of miRNA into mitochondria? We have observed an enhanced ER-mitochondria interaction upon HA-HuR expression to explain the increased import of miRNA into mitochondria (**Fig.4A-B**). To dissect the direct role of HuR in mitochondrial miRNA targeting, we isolated mitochondria from Huh7 cells for an in vitro miRNA targeting assay conducted in a reaction buffer containing 1 mM ATP. We noted a time-dependent entrapment of RNase-resistant miR-122 after 15 minutes of incubating the single-stranded synthetic miR-122 with mitochondria in a reaction buffer with 1 mM ATP and 250 mM KCl (**Fig. 4G and H**). Using mitochondria isolated from amino acid-fed and starved conditioned Huh7 cells, we detected a higher import of miRNA into the mitochondria isolated from starved Huh7 cells compared to those from fed control cell-derived mitochondria (**Fig. 4H**), possibly due to a greater amount of HuR associated with them. To confirm that a higher amount of HuR plays a role in miRNA import into mitochondria in vitro, we performed the miR-122 import reaction in the presence of 200 nM recombinant HuR (rHuR). We documented greater than two-fold enhancement in mitochondrial miR-122 content in the presence of HuR compared to a BSA Control (**Fig. 4I**). Thus, HuR can promote the entry of miR-122 into mitochondria under both *in vivo* and *in vitro* reaction conditions.

To explore the mechanism of HuR-mediated miRNA import into mitochondria, we investigated whether HuR remains bound to miRNAs inside mitochondria or if the ubiquitination of HuR, which plays a pivotal role in the endosomal targeting of miRNAs for EV-mediated export, is essential for their mitochondrial targeting. The inhibitor of EV-mediated miRNA export does not impact mitochondrial miRNA content; thus, miRNA targeting to mitochondria may not be coupled with EV-mediated export, a process controlled by ubiquitination and miRNA binding-unbinding of HuR (Mukherjee *et al*., 2016). We have used plasmids encoding full-length HA-HuR and HA-HuRΔH, a truncated mutant of HuR lacking the hinge region between RRMII and III, to transfect Huh7 cells. Unlike the full-length HuR, HuRΔH cannot be ubiquitinated and fails to promote miRNA export from hepatic cells due to the absence of potential ubiquitination sites in the deleted hinge region (Mukherjee *et al*., 2016). With an expression of HuRΔH, we did not observe any significant changes in the expression of Ucp2, Cytochrome C, or Mfn2 (**Fig.4**J and K). Both full-length and truncated HuR were found to associate with mitochondria (**Fig.4K**) under both fed and amino-acid-starved conditions. To evaluate the ubiquitination status of mitochondria-localized HuR, we isolated total and mitochondrial extracts to immunoprecipitate compartment-specific HuR using anti-HuR antibodies from control and HA-Ub expressing Huh7 cells. When western blotted with an HA-specific antibody, we detected ubiquitinated HuR only in the cytoplasmic extract of HA-Ub expressing Huh7 cells, while the mitochondrial extract predominantly contained non-ubiquitinated HuR (**Fig. 4L**). Consistent with HuR and HuRΔH both being associated with mitochondria, we observed enhanced levels of miR-122 and let-7g in starved Huh7 cell-derived mitochondria expressing HuRΔH (**Fig. 4M**). Through HA-immunoprecipitation of HA-HuR and HA-HuRΔH, we noted an increased association of miRNAs with HuR and its truncated version under starvation conditions (**Fig. 4N**). These data suggest that HuR-bound miRNAs remain stored in mitochondria during starvation and do not uncouple from HuR due to the absence of HuR ubiquitination, a process that is essential for endosome loading and the export of miRNAs via EVs. Consequently, compared to full-length HuR, which also promotes miRNA export via EVs in stressed hepatic cells, HuRΔH-unable to promote EV-mediated miRNA export-exerts a stronger effect on the mitochondrial localization of miRNAs and remains associated with mito-miRs in the mitochondria of starved Huh7 cells.

### Retro trafficking of mitochondrial miRNAs to the cytosol for its Ago re-association and formation of functional miRNPs upon amino-acid starvation-related stress reversal

We have observed the restoration of miR-122 activity in hepatic cells when they are refed with amino acids by placing them in amino acid-supplemented growth media (Bose & Bhattacharyya, 2016). We were interested in checking whether refeeding cells with amino acids might influence the mitochondrial miRNA content in refed hepatic cells. Refeeding the starved Huh7 cells resulted in shape, size, and volume changes. The increased size and volume of the mitochondria in starved Huh7 cells returned to control (fed) levels within 8 hours of refeeding (**Fig. 5A-B**). The elevated expression of the stress marker eIF-2α was also restored to control levels in refed Huh7 cells (**Fig. 5C**). We observed that the cellular miR-122 level, which drops in starved cells, was regained after refeeding with amino acids, while let-7a miRNA, which is not an abundant mitochondrial miRNA, showed an increase in its cellular level in starved cells and remained unaffected in refed cells (**Fig. 5D**). We noted increased levels of let-7g and miR-122 in the mitochondrial fraction only in starved cell-derived mitochondria, while these levels returned to control fed state in the mitochondria isolated from amino acid-refed cells (**Fig. 5E**). Thus, the mitochondrial and total cellular levels of miRNAs change in opposite directions, suggesting possible physiological implications of mito-miR’s storage in mitochondria.

**Figure 5.**
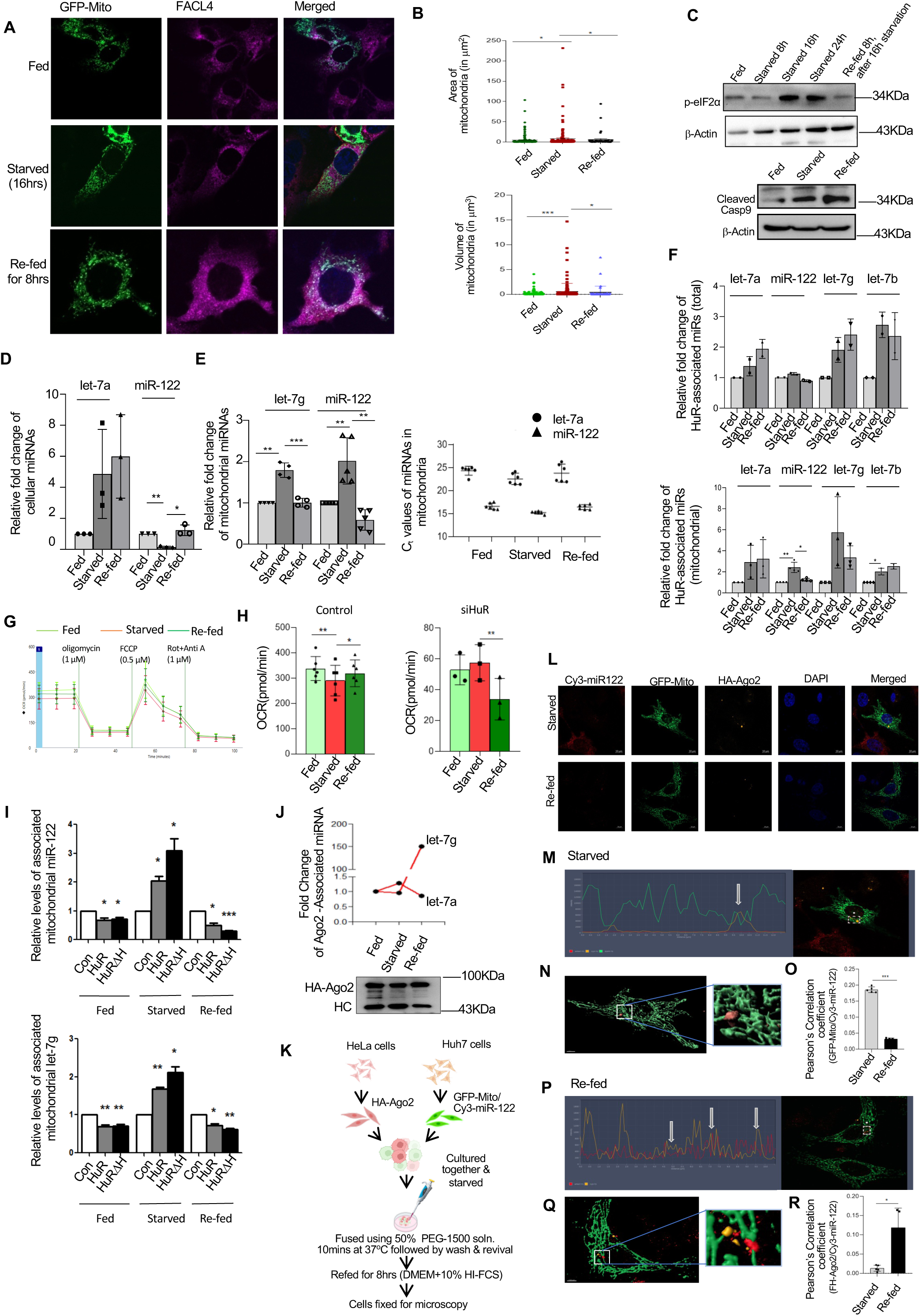
Reversing starvation-induced stress is linked to the re-localization of mitochondria to the cytosol and the re-association of Ago2 with mito-miRs. A-B. Effect of amino acid starvation and refeeding of Huh7 cells on mitochondrial shape (A), size, and volume (B). All experiments were conducted in triplicates. **C.** Expression of the stress marker phosphorylated eIF-2α and the apoptosis-related marker cleaved PARP in fed, starved, and re-fed Huh7 cells. A Western blot of the respective proteins was conducted using cell lysates obtained from Huh7 cells subjected to fed, starved, and re-fed conditions. **D.** Relative changes in cellular miRNA content in Huh7 cells were subjected to feeding or 16 hours of amino acid starvation, followed by 8 hours of re-feeding after the starvation period. Values were normalized against U6 RNA for quantification. **E.** Changes in mitochondria-associated miRNAs in Huh7 cells subjected to amino acid-fed, starved, and re-fed conditions are described above. Relative values are plotted with the fed condition value set as a unit. The values are normalized against let-7a miRNA levels. Changes in Ct values for let-7a and miR-122 are plotted in the right panel. All experiments were conducted in triplicates. **F.** Relative changes in HuR-associated miRNA levels in cytosolic and mitochondria-associated HuR of Huh7 cells under amino acid-fed, starved, and re-fed conditions. Values are derived from three independent experiments and normalized against the amount of HuR that was immunoprecipitated. **G.** Oxygen consumption rate changes with amino acid starvation and is restored upon refeeding. The effect of starvation and refeeding on the oxygen consumption rate (OCR) of Huh7 cells subjected to amino acid starvation and refeeding is plotted against the time of injections of oxidative phosphorylation and electron transport chain blockers. **H.** Change in the oxygen consumption rate (OCR) of control and siHuR-treated Huh7 cells under fed, starved, and re-fed conditions. The data confirmed impaired OCR in HuR-depleted re-fed cells, suggesting a lack of respiratory fitness during the recovery from starvation, which may account for the accelerated cell death and reduced OCR in HuR-depleted Huh7 cells in the re-fed condition. **I.** HuR and HuRΔH demonstrate the reversibility of their bindings with mito-miRs in amino acid-fed, starved, and re-fed cells. Mitochondrial HuR and HuRΔH exhibit a substantial increase in miRNA binding under starvation conditions, which reverses in re-fed cells. Values are derived from three independent experiments and normalized against the amount of HuR immunoprecipitated. **J.** Reloading mitochondria-relocalized cytoplasmic miRNAs with Ago2 in stress-reverted hepatic cells. The association of let-7g and let-7a changes with immunoisolated HA-Ago2 is analyzed in fed, starved, and re-fed conditions. The western blot of FH-Ago2 is displayed at the bottom, and the band intensity of Ago2 is used to normalize miRNA quantity. **K.** This is a scheme for an experiment involving the fusion of HeLa and Huh7 cells that express HA-Ago2 and GFP-Mito expression plasmids, respectively. The Huh7 cells were transfected with Cy3-mi-122, and both cell types were cocultured before being starved and subsequently fused with PEG-1500, before being refed with amino acids. **L.** Representative microscopy panels show cells after 16 hours of starvation or 8 hours of refeeding following starvation. GFP-tagged mitochondria are shown in green, Cy3-tagged miR-122 in red, HA-Ago2 in far red (yellow), and DAPI stains the nucleus (blue). The merged panels appear on the right **M.** Graphs illustrate the relative pixel density of Cy3-tagged miR-122 compared to Mito-GFP for regions highlighted within the inset frames of the image in panel L. White arrows indicate overlapping pixel intensity peaks, signifying the co-localization of the two entities **N.** 3D-rendered model of mitochondria in Mito-GFP expressing Huh7 cells transfected with Cy3-miR-122 after 16 hours of starvation and its fusion with HA-Ago2 HeLa cells. It illustrates co-localized Cy3-tagged miR-122 on the surface of the mitochondrial tubules, with little or no association with Ago2. The marked area is zoomed in to display the colocalization in 3D. **O.** Graph illustrating Pearson’s coefficients for the co-localization of Cy3-miR-122 and GFP-Mito in fused cells under starved or refed conditions. **P.** Graphs depicting the relative pixel density of Cy3-tagged miR-122 compared to HA-Ago2 for regions highlighted within the inset frames taken from panel L. White arrows indicate overlapping pixel intensity peaks, signifying the co-localization of the two entities. **Q.** The 3D-rendered model of mitochondria shows co-localized Cy3-tagged miR-122 with HA-Ago2 under re-fed conditions. **R.** Graph showing Pearson’s coefficients for co-localization of Cy3-miR-122 and HA-Ago2 in fused cells during starvation or refeeding conditions. Scale bar: 10 µm. Statistical significance was derived from the Student’s t-test. The mean ± s.e.m. is shown for n>3. *P<0.05, **P<0.01, ***P<0.001 (Student’s t-test).

We have documented an increased association of HuR with mitochondrial miRNAs, specifically miR-122 and let-7g, which reversed upon stress relief in re-fed cells. The total cellular miRNAs’ association with HuR remained high even after re-feeding the cells with amino acids (**Fig.5F**). We hypothesize that the HuR-driven mitochondrial localization of mito-miRs has significant implications for mitochondrial function. The oxygen consumption rate is a strong marker of cell fitness in response to challenges affecting oxidative phosphorylation (Chatterjee *et al*., 2020; Smolina *et al*, 2017). The HuR-driven miRNA localization, as impaired in siHuR-treated cells, was found to influence OCR. Upon depletion of HuR by siRNA in fed, starved, and re-fed Huh7 cells, we observe a significant decrease in OCR (oxygen consumption rate) in re-fed cells compared to the control non-siHuR treated cells, where a drop in OCR during starvation was recovered in the re-fed condition. This data suggests major mitochondrial dysfunction during stress reversal in cells depleted of HuR. Thus, the re-fed cells fail to regain proper function upon stress removal. This indicates that HuR-driven miRNA localization to mitochondria may be a key process in protecting amino acid-starved cells under metabolic stress and its reversal.

Interestingly, supporting the idea that HuR is crucial for the mitochondrial storage of miRNAs, we found that the HuR mutant HA-HuRΔH demonstrated a stronger miRNA binding-unbinding capacity for miR-122 and let-7g in starved and refed cell-derived mitochondria compared to wild-type HA-HuR (**Fig 5I**). Interestingly, while the association of let-7g miRNAs with HuR and its mutant version in mitochondria decreased in re-fed cells, the association of let-7g exhibited a markedly different pattern compared to its association with Ago2. Ago2 was found to associate more with let-7g in re-fed cells, whereas there was a decline under starved conditions (**Fig.5J**).

Could miRNAs that have been relocalized to the mitochondria associate with Ago2 to form de novo miRNPs in re-fed cells? To demonstrate that miRNAs relocalized from mitochondria are functional and associated with Ago2 in re-fed cells, we adopted a cell fusion assay to confirm the re-loading of miRNAs delocalized from mitochondria with Ago2 in re-fed cells (Yang & Shen, 2006) (**Fig. S4**). Using FH-Ago2-expressing HeLa cells (which do not express miR-122) and fusing them with Cy-miR-122-transfected and Mito-GFP-expressing Huh7 cells after starvation, we observed the recovery of the fused cells under amino acid re-feeding conditions. Notably, we detected FH-Ago2 association with Cy-3 miR-122 after the fusion of the cells in the re-fed state. In contrast, the association of Cy-miR-122 with mitochondria decreased (**Fig. 5 K-R**). This data suggests the re-association of Ago2 with mitochondria-delocalized miRNAs in re-fed cells.

### Mito-miRs specifically target mRNAs that encode mitochondrial proteins involved in apoptosis pathways

With amino acid starvation-induced stress reversal upon refeeding cells with amino acids, we detected polysome enrichment of miRNAs that relocated from mitochondria during refeeding. The targets of these miRNAs were found to be dislodged from polysomes, suggesting their effective translational repression during the refeeding phase (**Fig.6A-B**). This data supports a reversible expression regulation by mito-miRs occurring during the starvation and refeeding cycle. The transport of mito-miRs to mitochondria during starvation facilitates the expression of target mRNAs, including BCL2 and BCL2L1, which regulate cellular apoptosis. Once the cells reverted to the re-fed condition, the rapid re-localization of mito-miRs to the cytoplasm likely permits the re-repression of target mRNAs, thereby repressing the mRNAs encoding apoptosis and preventing cell death (**Fig.6A-B**).

**Figure 6.**
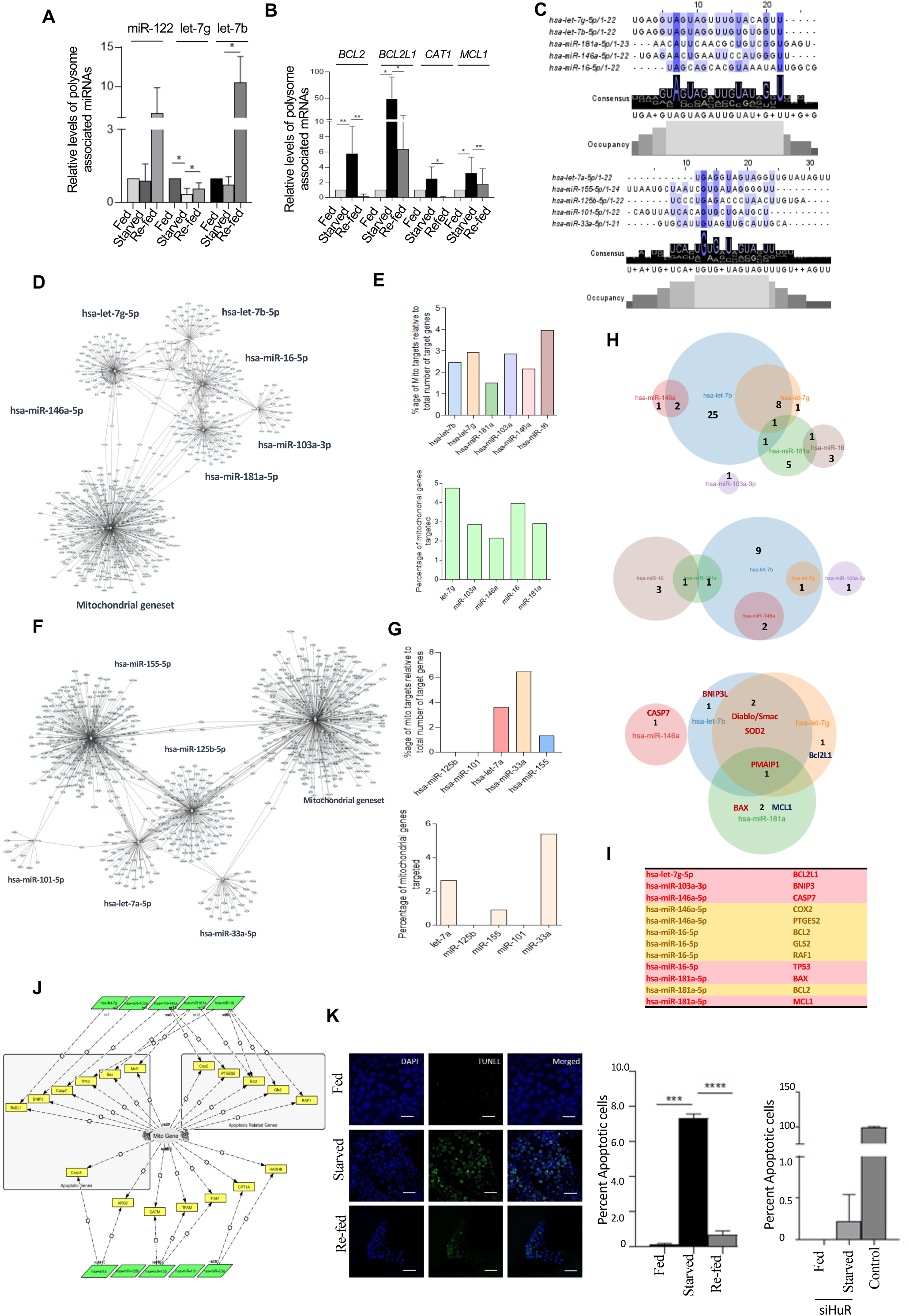
Mito-miR targets mRNAs encoding mitochondrial components of apoptosis-related pathways. **A.** The effect of re-feeding amino acid-starved cells on the polysome association of miRNAs was examined. Polysomes from fed, starved, and re-fed cells were isolated to quantify the miRNAs associated with the polysomes. The relative levels of mito-miR association change with starvation and are revised in re-fed cells. RNA was quantified by qRT-PCR and normalized against let-7a. Values from fed cell polysomes are considered units for each miRNA measured. **B.** Starvation-induced polysome association of genes are the known targets of mito-miRs quantified in RNA isolated from polysomes of fed, starved, and re-fed cells. The values are normalized against GAPDH mRNA levels, and the values obtained from fed cell polysomes are considered units. **C.** The mito-miRs and non-mito-miRs exhibit sequence variability and consensus among themselves. A sequence comparison using MAFFT algorithm-based software determines the percentage similarity of the mito-miRs and non-mito-miRs. **D.** A comprehensive network depicting differential mitochondrial miRNA distribution alongside a complete set of mitochondrial genes mapped to a network of mito-miR targets. Nodes within the network signify each gene involved in the interactions. At the same time, edges denote instances of validated gene targets for each miRNA (redundant targets represented by multiple nodes correspond to several validation instances identified through data mining from the available literature. **E.** Graphical representation of the percentage of mitochondrial geneset mRNA targets regulated by individual mito-miRs relative to the total number of target mRNAs known to be regulated (as represented in the network) (top panel). Graphical representation of the percentage of mitochondrial geneset targets regulated by individual mito-miRs (bottom) panel). **F.** A network representing the differential distribution of non-mitochondrial miRNAs and a complete mitochondrial gene set is mapped to a network of non-mito-miR targets. The nodes in the network represent each gene involved in the interactions, while the edges indicate validated gene targets for each miRNA (redundant targets with multiple nodes signify several instances of validation obtained from data mining available in papers and texts) **G.** Graphical representation of the percentage of mitochondrial geneset targets regulated by individual non-mito-miRs relative to the total number of targets they regulate (as shown in the network) (top panel). Graphical representation of the percentage of mitochondrial geneset targets regulated by individual non-mito-miRs (bottom panel) panel). **H.** Venn diagrams illustrate the number of individual targets and shared ones that correspond to the mitochondrial geneset stated here, as determined through network analysis (top panel). Venn diagrams showcase the total number of unique and shared mitochondrial component-encoding genes associated with the apoptotic pathway and targeted by the mito-miRs (middle panel). Venn diagrams represent the genes regulated by mito-miR that are members of the apoptotic pathway (bottom) panel). **I.** Mitochondrial gene set members (genes encoding proteins involved in mitochondrial functions) targeted by mito-miRs. **J.** The diagram illustrates the targets of mito-miRs and non-mito-miRs that can be mapped to the mitochondrial gene set, which was constructed as a small network. Apoptotic pathway genes and related genes that these miRNAs target are clearly indicated within defined boxes featuring various shades. Function arrows point to nodes representing different mRNAs/genes, with the arrows denoting the roles of these targets in relation to miRNAs. Apoptosis is induced in fractions of cells by starvation and returns to normal levels under re-fed conditions, as determined by the TUNEL assay performed on respective amino acid-fed, starved, and re-fed Huh7 cells. The effect of siRNA-mediated knockdown of HuR on starvation-induced apoptosis is illustrated in the right panel. A positive control was established using DNase-treated cells. Scale bar: 10 µm. Statistical significance was determined using the Student’s t-test. Values are shown as mean ± s.e.m. for n>3. *P<0.05, **P<0.01, ***P<0.001 (Student’s t-test).

We were interested in whether the localization and relocalization of mitochondria-localized mito-miRs contribute to stress-induced changes in mRNA expression in mammalian cells. Understanding that miRNA storage is a defense mechanism against cellular stress, we explored whether mito-miRs specifically regulate a subset of mRNAs during the stress response and refeeding phase. Our aim was to determine whether mito-miRs preferentially target genes encoding proteins related to apoptosis and stress response. Mito-miRs clearly have a sequence consensus that differs from that of the non-mitochondrial miRNA cluster (**Fig. 6C**). We analyzed the mRNAs targeted by mitochondrially located mito-miRs. We compared them with those targeted by non-mitochondrial miRNAs. We noted that mitochondrially located mito-miRs preferentially target genes encoding mitochondrial proteins. Through a dataset-based miRNA analysis of the global gene set versus the mitochondrial protein-coding gene set, we observed a clear preference for individual mito-miRs targeting mitochondrial protein-coding nuclear genome-encoded genes compared to what we observed with non-mitochondrial miRNAs (**Fig. 6 D-G**). Among the mitochondrial gene set, mito-miRs regulate a higher percentage of mitochondrial component-encoding genes that are targeted by non-mitochondrial miRNAs. Mito-miRs target 14.29% of total mitochondrial genes. In contrast, only a small fraction of the mitochondrial genes are regulated by individual non-mitochondrial miRNAs (**Fig. 6 F-G**).

What implications does this observation have for cellular physiology? While examining the gene set regulated by mitochondrial miRNAs, we found that mitochondrially located mito-miRs target a significant portion of mitochondrial genes encoding apoptosis-related pathways. In contrast, in the analysis conducted for non-mitochondrial miRNAs, we identified that only a small number of apoptosis-related genes are targeted by non-mitochondrial miRNAs (**Fig. 6 H-J and Fig. S 5**). This suggests that mitochondrial miRNAs are imported into mitochondria under stress conditions to enable the activation of apoptosis-related proteins, inducing apoptosis in starved cells, which can be reversed in re-fed cells, possibly due to the relocalization of miRNAs from the mitochondrial matrix to the cytoplasm and subsequent reactivation of Ago-miRNPs formed by relocalized mito-miRs in the cytoplasm of the re-fed cell.

We conducted target analysis to identify gene sets regulated by specific groups of miRNAs and are already predicted or validated targets of the respective miRNAs. We detected that mitochondrial miRNAs target more components of apoptosis pathways and related targets than non-mitochondrial miRNAs in the human gene set (**Fig. 6 J**). miR-122, which also shows shuttling to and from mitochondria during the starvation-refeeding cycle, targets three apoptosis and related pathway genes, namely MCL 1, Bax, and BCl 2 l 1, suggesting a similar regulation of apoptosis by miR-122 as occurs with other mito-miRs in hepatic cells. Confirming the upregulated expression of apoptosis-related genes and the association of encoding mRNAs with polysomes (**Fig. 6 A-B**), we noted enhanced apoptosis in the starved cell population, followed by a reversal to fed levels in re-fed cells (**Fig. 6 K).** Depleting HuR using siHuR, which prevents miRNA localization to mitochondria, caused reduced starvation-induced apoptosis in Huh 7 cells. This signifies the importance of cytosolic mito-miRs availability for stress-induced apoptosis (**Fig.6K**).

Like our observations with humans, we found similar data when analyzing murine gene sets for mitochondrial miRNAs. Among the murine target genes related to mito-miRs, a significant portion of the gene set encoding mitochondrial components, which is regulated by mitochondria-localized mRNAs or mito-miRs, was found to be linked with apoptosis pathways (**Fig. S6**).

## Discussion

Notably, the targeting of nuclear-encoded gene products from eukaryotes to mitochondria, such as 5S RNA and tRNAs, has been documented throughout the evolutionary process. (Bhattacharyya & Adhya, 2004; Salinas *et al*, 2008; Smirnov *et al*, 2011; Warren *et al*, 2021). Unicellular protozoan parasites lack the genes that encode tRNAs in their mitochondrial genomes, instead importing cytosolic tRNAs to facilitate the translation of mRNAs encoded by the mitochondrial genome (Bhattacharyya & Adhya, 2004; Salinas *et al*., 2008; Warren *et al*., 2021). The presence of miRNAs within mitochondria has been observed in various eukaryotic systems; however, the implications of this localization remain unclear (Canale & Borghini, 2024; Monzel *et al*, 2023; Skawratananond *et al*, 2025). Recent reports suggest that miRNAs may be targeted to mitochondria to inhibit genes encoded by mitochondria (Luo *et al*., 2024; Roman *et al*, 2020). We and others have observed the lack of Ago proteins in purified mitochondria from human cells that are free of endoplasmic reticulum (ER) and MAM contamination (Maniataki & Mourelatos, 2005). This suggests a potential uncoupling of Ago2 from miRNAs located in mitochondria; thus, mitochondria-enriched miRNAs may not perform their usual gene-repressive functions within the mitochondrial context due to the lack of Ago proteins in mitochondria. However, subcellular localization studies with other Ago protein isoforms in human or murine cells are lacking, making it impossible to conclude their absence in mammalian mitochondria.

What roles do Ago-uncoupled miRNAs play within mitochondria? Our observations reveal that when human hepatocytes encounter metabolic stress, the localization of miRNAs in mitochondria increases significantly. A similar phenomenon likely occurs when cells face other forms of stress. Remarkably, numerous genes require upregulation during stressful conditions, many of which are regulated by miRNAs (Leung & Sharp, 2010). Consequently, an intriguing hypothesis suggests an uncoupling of miRNAs from Ago2 and their subsequent import into mitochondria—potentially in an Ago2-uncoupled state—as a storage mechanism for inactive miRNAs intended for future use. The fate of these mitochondrial miRNAs is also fascinating, as the miRNA storage mechanism seems reversible. When stress responses are alleviated, genes upregulated due to stress must be re-repressed again. Mitochondrial “stored miRNAs” return to the cytoplasm and re-engage with the Ago2 pool upon stress reversal. Various factors could influence the fate of mitochondrial miRNAs. Recent reports suggest the selective export of these miRNAs via mitovesicles that also carry components of the mitochondrial respiratory complex (D’Acunzo *et al*, 2024; D’Acunzo *et al*, 2022; D’Acunzo *et al*, 2021). Mitophagy remains a crucial mechanism for decreasing mitochondrial RNA, thereby protecting cells from dysfunctional mitochondria and malformed organelles (Harbauer *et al*, 2022). However, we have not observed any impairment; instead, there is an increase in mitophagy in cells experiencing starvation-related stress, which does not account for the elevated miRNA content within the mitochondria seen in stressed cells.

The mechanism of miRNA storage within mitochondria may also occur in other types of mammalian cells, ensuring the effective execution of its dual functions. In addition to buffering cellular miRNA activity and promoting the translational upregulation of mRNAs that are inhibited by miRNAs in human cells, another potential role of mitochondrial targeting of miRNAs under stress involves regulating miRNA export and mitigating the dissemination of EV-mediated signals to neighboring cells, thereby potentially restoring inflammatory responses in those cells (Duroux-Richard *et al*., 2021; Ma *et al*., 2023a). This indicates a vital mechanism for safeguarding affected tissues from the dissemination of inflammatory signals by sequestering inflammatory miRNAs, such as miR-122, in the mitochondria of hepatic cells within an in vivo context, such as during the starvation of mice for amino acids or exposure to an MCD diet (Bandyopadhyay *et al*., 2023).

Our research indicates that mitochondrial miRNA targets nuclear-encoded mRNAs that preferentially encode mitochondrial components. This observation is particularly compelling. Previous studies suggest that eukaryotic ribosomes may remain associated with the outer membrane of mitochondria, facilitating the engagement of mRNAs encoding mitochondrial proteins with mitochondrial membrane-associated polysomes (Lesnik *et al*, 2015). This may explain the highly regulated translational response seen in cells undergoing metabolic changes that necessitate swift alterations in mitochondrial membrane potential. We propose the hypothesis that mitochondrial-targeted miRNAs may aid in the recruitment of these target mRNAs to the mitochondrial surface, where they attach to polysomes for a timely and context-dependent fast and tunable response required for eukaryotic cells in fluctuating environments. Our data suggest that a pool of Ago2-bound mitochondrial miRNAs accumulates around the mitochondria and may act as an anchor for mRNAs to be recruited to polysomes associated with mitochondria. Additionally, our findings support the idea that the mitochondrial localization of miRNAs is reversible, allowing for quick adjustments of miRNA repression on target mRNAs associated with polysomes located near the mitochondria or attached to its outer membrane.

The regulation of miRNA storage within mitochondria is influenced by RNA-binding proteins such as HuR. Intriguing evidence suggests that HuR-miRNA binding is essential for the entry of miRNAs into mitochondria. Reports indicate that the HuR-miRNA complex can also be targeted to endosomes (Goswami *et al*., 2020; Mukherjee *et al*., 2016). The interactions between mitochondria and the endoplasmic reticulum regulate endosomal miRNA levels as well as the miRNA recycling and export process (Bose *et al*., 2020; Chakrabarty & Bhattacharyya, 2017). However, interactions between ER and mitochondria are compromised by the protozoan parasite *Leishmania donovani*, which impairs the endosomal targeting of miRNAs and Ago2 (Chakrabarty & Bhattacharyya, 2017). In macrophages infected with Leishmania donovani, HuR is notably cleaved, hindering miRNA export (Goswami *et al*., 2020). Intriguingly, HuR, which lacks a domain necessary for ubiquitination, does not disrupt its binding with miRNAs but does affect its export from mammalian cells via EVs (Mukherjee *et al*., 2016). When ectopically expressed in hepatocytes, this mutant increased the loading of mitochondrial miRNAs, while the same miRNAs underwent restricted EV-mediated export (Mukherjee *et al*., 2016). This indicates that HuR ubiquitination serves as a crucial signal that determines the distribution of HuR-miRNA complexes, promoting either miRNA storage or export. Our observations reveal a unique function of miRNA storage within mammalian mitochondria in an Ago2-uncoupled form. We have documented that impairing miRNA storage in mitochondria leads to reduced apoptotic pathways in cells associated with compromised mitochondrial oxygen consumption. The reversible and contextual storage of miRNAs in mitochondria may represent a widely adopted mechanism in mammalian somatic and neuronal cells for reversing target gene expression in response to stress or external stimuli, such as neurotransmitters, and for balancing cell death and survival excitability.

Interestingly, our data suggest that inhibition of extracellular vesicle (EV) release does not affect the levels of certain mitochondrial miRNAs (e.g., miR-122), but it does increase their cellular accumulation (Fig. 3M). Under starvation conditions, we observed a significant decrease in cellular levels of some non-mitochondrial miRNAs (e.g., let-7a), while their mitochondrial levels remained unchanged. This indicates potential EV-mediated export as a primary mechanism for non-mitochondrial miRNA regulation, as starvation may induce the primary ciliogenesis process that could enhance EV production release.

Recent research shows a significant increase in these mitochondrial-enriched miRNAs, known as mito-miRs, in extracellular vehicles (EVs) released by morphine-treated astrocytes (Ma *et al*, 2021; Ma *et al*, 2023b). These miRNAs play a crucial role in promoting ciliogenesis in naive astroglial cells (Ma *et al*., 2021; Ma *et al*., 2023b). We propose that targeting mitochondrial miRNAs in morphine-treated astroglial cells promotes their release through mitovesicles, which are unique, mitochondria-derived vesicles released by morphine-treated astroglia. We suggest that the initial targeting of these miRNAs to mitochondria in animal cells depends on the interplay between mitochondria and the endoplasmic reticulum (ER), a process further enhanced by the RNA-binding protein ELAVL1 (HuR), which regulates miRNA activity and export across various mammalian cell types.

However, several aspects of mitochondrial storage of miRNAs remain unexplored. Since the miRNAs within the mitochondria show no association with Ago, as minimal Ago2 is found in pure mitochondria devoid of any ER and MAM contamination, it is crucial to understand how mitochondrial miRNAs function as target mRNA repressors within the mitochondria. Can these mito-miRs possibly act as part of the HuR-miRNA complex to regulate mRNA stability provided by HuR? Given that both HuR and HuR-associated miRNAs are prevalent in mitochondria from amino acid-starved Huh7 cells, it may be that the mito-miR recruits HuR to those target mRNAs to stabilize them within the mitochondria.

Recent reports indicate that mitochondria can be transported from one cell to another through tubular structures that enable the exchange of mitochondrial content between cells linked by intercellular tubular tunnels or tunneling nanotubes (TNT) (Wang & Gerdes, 2015). Mitochondrial transport has also been observed along the dendrites of neurons, connecting their proximal and distal ends. Here, selective mRNAs are translated locally in response to external signals and the effects of neurotransmission at both postsynaptic and presynaptic sites (Sheng & Cai, 2012; Zinsmaier *et al*, 2009). In both cases mentioned above, it is possible that mito-miRs are transported along with the mitochondria and released locally to influence the translation of proteins in distant locations.

## Materials and Methods

### Cell Culture, starvation, refeeding, and transfections

Human HeLa, Huh7, mouse embryonic fibroblast (MEF-all genotypes) cells were cultured in Dulbecco’s Modified Eagle’s medium (DMEM; Gibco-BRL) supplemented with 2mM L-glutamine and 10% heat-inactivated fetal bovine serum (FBS). MEF-WT and Mfn2 ^-/-^ MEF cells were obtained from ATCC. All cell culture reagents were obtained from Life Technologies, USA. For starvation experiments, cells were treated with a medium comprising Hank’s balanced salt solution (HBSS) and 10% dialyzed fetal bovine serum (FCS) for 16 hours while the control fed or re-fed medium contained Hank’s balanced salt solution (HBSS) and 10% heat-inactivated non-dialyzed fetal bovine serum (FCS) was used as described before (Bhattacharyya *et al*., 2006; Mukherjee *et al*., 2016). Re-feeding was done after 16 hours of starvation. Following the manufacturer’s instruction, plasmid transfections were done with Lipofectamine 2000 (Life technologies) in Opti-MEM media. Indicated cell types in various experiments were treated with Thalpsigargin (Sigma, 2.5 µM) or H_2_O_2_ (3mM) and Tunicamycin (Sigma, 50 µg/ml). Seahorse oxygen consumption experiments were done using all kit-supplied reagents from Agilent technologies. This was done in Seahorse equipment (Seahorse Bioscience, North Billerica, MA, USA.) essentially as described previously with Fed, Starved, and Re-fed cells(Bose *et al*., 2020; Chatterjee *et al*., 2020).

### siRNA and plasmid transfections

SiRNAs against different proteins were purchased from Dharmacon. All Dharmacon SMART Pool ON-TARGET plus were procured. According to the manufacturer’s instructions, siRNA transfection was done using RNAi Max (Life Technologies). Huh7 cells were transfected with 25 moles of siRNA per well of a 24-well plate. The siRNA-transfected cells were split 24 hrs later and were harvested 72-96 hours post-transfection to ensure proper knockdown of the siRNA target gene. All plasmid transfections were done for 6hrs followed by 48h incubation after splitting of cells done after 24h of transfection to promote transient expression of the specific proteins. Details of plasmids are available in Supplementary Table S1.

### Animal experiments

BALB/c mice of around 4-6 weeks of age were maintained in two groups of 3 animals each for the different conditions of fed and starved animals. Fed state or control animals were provided with a normal chow diet (pellets containing amino acids, carbohydrates, and normal concentrations of salt) and water. The amino acid starvation was administered to animals by feeding them with sugar cubes and saline water for 16 hrs (overnight). On the next day, animals were sacrificed, and liver samples were collected in PBS and later kept in an isolation buffer supplemented with 0.5% BSA. For MCD diet-induced changes in mitochondrial miRNA levels in murine hepatic cells, Eight- to ten-week-old male C57BL/6 mice (20–24 g) were housed in controlled conditions (23 ± 2 °C, 12 h light/dark cycle) in individually ventilated cages. They were randomly divided into two groups and fed either a standard chow diet or an MCD diet (MP Biomedicals; catalog no. 0296043910) for 4 weeks. The Animal Ethics Committee of CSIR-Indian Institute of Chemical Biology, India, approved the use of mice for experiments. Experiments during the study were carried out in accordance with the National Regulatory Guidelines issued by the Committee to Supervise Experiments on animals, Ministry of Environment and Forest, Government of India.

### MAM isolation procedure

Isolation of mitochondria and mitochondria-associated membrane (MAM) in our study has been generally done from cells gathered from 70-80% confluent 3 to 5 cell culture dishes of 100 mm diameter. The cells from the culture dishes were scrapped and collected in DPBS, which lacked calcium and magnesium ions (-Ca^2+^, -Mg^2+^) and was not done using the general trypsinization method. Then, these cells were centrifuged twice at 600xg for 5 mins at 4°C in DPBS. All further steps involving centrifugation were done at 4°C. The cells were collected, and the supernatant was discarded during these centrifugation steps. The cell was then resuspended in isolation buffer-1 (IB-1), and samples were rotated for 30 minutes on a rotator. After 30 minutes of incubation, the IB-1 cells were collected in a pre-cooled cell homogenizer, and approximately 70 strokes were given per sample for lysing the hepatic HUH7 or Hep-G2 cells (50 strokes in the case of HeLa and MEF cells). The lysate thus obtained from the homogenizer was then centrifuged at 600xg for 5 mins twice. The sup was collected in each case, and the cellular debris was obtained as the pellet was discarded. This sup was then subjected to another round of centrifugation at 7000xg for 10 mins and the sup thus obtained was collected in another microcentrifuge tube. This sup contained crude fractions of ER along with lysosomal contaminations. The pellet containing crude mitochondrial extracts was resuspended in another isolation buffer (IB-2), and the fractions were again centrifuged at 7000xg for 10 mins, and the sup obtained was again discarded. Obtained pellets were resuspended in IB-2 and centrifuged at 10,000xg for 10 minutes and the final crude mitochondrial pellet thus obtained was resuspended in mitochondria resuspension buffer (MRB). A basal medium was prepared and then mixed with Percoll^TM^ to form a 30 percent personal cushion from which 8 ml was loaded into a centrifuge tube. The mitochondrial lysate in MRB buffer was loaded on top of this cushion. The rest of the tube was carefully filled with MRB buffer on top of the lysate. This tube was then centrifuged in a Beckman Coulter ultracentrifuge at 95,000xg for 30 mins, and the pure mitochondrial pellet was obtained on the Percoll^TM^ collected at the bottom of the tube. MAM region appears as a translucent floating ring somewhere in the center of the tube, and both these fractions were collected with loading tips with the help of micro-pipettes. The MAM fraction is diluted to ∼10 times in MRB and is subjected to centrifugation at 100,000xg for 1 hr. The crude ER fraction that was collected earlier during crude mitochondrial fractionation was also centrifuged at 20,000xg. The supernatant was collected and further run on the ultracentrifuge alongside the MAM fractions for 1 hr at 100,000g. At the end of this extensive centrifugation process, MAM is obtained at the bottom of the centrifuge tube as a floating pellet, and the ER is obtained similarly but as a relatively harder pellet. The fractions were then collected individually and processed for protein estimation or RNA isolation with Trizol LS reagent.

In the case of MAM isolation from animal liver samples, the isolation buffers 1 and 2 were supplemented with 0.5% bovine serum albumin (SRL). The lysis of the liver samples was achieved by using a mechanical tissue homogenizer, keeping the speed of the rotor at 3,000-4,000 rpm to lyse the cells without affecting the structure of the mitochondria and other organelles.

The IB-1 contains 225 mM mannitol, 75 mM sucrose, 0.1 mM EGTA, and 30 mM Tris-HCl, pH 7.4. IB-2 has the same composition as IB-1 but without EGTA. The MRB (mitochondria resuspending buffer) contains 250 mM mannitol, 5 mM HEPES (pH 7.4), and 0.5 mM EGTA. The Percoll medium was of the composition of 225 mM mannitol, 25 mM HEPES (pH 7.4), 1 mM EGTA, and 30% Percoll (vol/vol).

### Measurement of Mitochondrial Oxygen Consumption

Huh7 cells were plated accordingly as fed, starved or refed cultures in 24 well XF-cell plates (Seahorse bioscience) in 100µl DMEM growth medium containing 10% FBS and placed in a 5% CO2 incubator. Once the cells adhered (4-5hrs later), 150µl growth medium (containing normal or Dialyzed FBS) was added into each well, and the cells were maintained overnight at 37^0^C in a 5% CO_2_ incubator. On the following day, refeeding was carried out as per our treatment protocols, and all time points were adhered to for further workflow. DMEM base medium (Seahorse Bioscience, North Billerica, MA, USA) supplemented with 25µM D-glucose and 1µM sodium pyruvate (adjusted to pH 7.4) was added to the cells and incubated for 1 hr in a non-CO2 incubator at 37^0^C. The three injection ports (A-C) of the XFe cartridge were loaded with Oligomycin (Oligo, 1μM), Carbonyl cyanide-4-trifluoromethoxyphenylhydrazone (FCCP, 0.5 μM) and Rotenone and antimycin A (Rot + Anti A, 1μM) respectively followed by equilibration and calibration in the instrument for 12 min. Following this, the cell plate was loaded to initiate measurement of oxygen consumption rate (OCR) following the 3 min wait, 2 min mix and 3 min measurement cycle over a total period of approx. 1hr.

Protein estimation was done by ELISA using Bradford’s reagent to normalize the obtained OCR values OCR rate which was expressed in pmol/min/mg protein. The overall data was analyzed in Wave software, which is available on the Agilent Technologies website as a platform for XFe analyzer tests. Normalized OCR data usually indicates the levels of mitochondrial respiration exclusively and rules out any other probable interfering sources of oxygen consumption.

### Western blotting

The samples were diluted in 5X sample loading buffer (312.5 mM Tris-HCl pH 6.8, 10% SDS, 50% glycerol, 250 mM DTT, 0.5% bromophenol blue) and heated for 10 minutes at 95°C. Following SDS-polyacrylamide gel electrophoresis, proteins were transferred to PVDF nylon membranes. Membranes were blocked in TBS (Tris-buffered saline) containing 0.1% Tween-20 and 3% BSA. Primary antibodies were added in 3% BSA for a minimum 16 hrs at 4°C. Following overnight incubation with antibody, the membranes were washed at room temperature thrice for 5 min with TBS containing 0.1% Tween-20. Washed membranes were incubated at room temperature for 1 hr with secondary antibodies conjugated with horseradish peroxidase (1:8000 dilutions). Excess antibodies were washed three times with TBS-Tween-20 at room temperature. Antigen-antibody complexes were detected with West Pico Chemiluminescent or West Femto Maximum Sensitivity substrates using standard manufacturer’s protocol (Thermo-Scientific). Imaging of all western blots was performed using an UVP Bio-Imager 600 system equipped with VisionWorks Life Science software (UVP) V6.80. Antibodies used in this study are listed in Supplementary Table 2.

### RNA Isolation and Real-Time PCR

According to the manufacturer’s protocol, total RNA was isolated using TriZol or TriZol LS reagent (Invitrogen). MiRNA assays by real-time PCR were performed using specific primers for human let-7a (assay ID 000377), human miR-122 (assay ID 000445), human miR-21 (assay ID 000397), human miR-181a (assay ID 000480), human let-7b (assay ID 000378), human let-7g (assay ID 002282). U6 snRNA (assay ID 001973) was used as an endogenous control. Real-time analyses by two-step RT-PCR were performed for quantification of miRNA levels on Bio-Rad CFX96TM real-time system using Applied Biosystems TaqMan chemistry-based miRNA assay system. One-third of the reverse transcription mix was subjected to PCR amplification with TaqMan Universal PCR Master Mix No AmpErase (Applied Biosystems) and the respective TaqMan reagents for target miRNA. Samples were analyzed in triplicates. The comparative C_t_ method, which included normalization by the U6 snRNA was used for relative quantification. To quantify mRNA, we prepared cDNA from RNA samples using the Eurogentec Reverse Transcriptase Core Kit. Real-time PCR was conducted on the cDNA utilizing the Mesa Green qPCR Mastermix Plus for SYBR Assay-Low ROX (Eurogentec). We used 18 s rRNA as an endogenous control.

### Polysome isolation

For polysome isolation, approximately 1 × 10^7^ cells were used as starting material. Cells were collected by scraping in PBS and initially were centrifuged at 600xg for 5 mins at 4°C. The cell pellet was then incubated (kept in rotation) in 1X lysis buffer (10 mM Hepes, 25 mM KCL, 5 mM MgCl_2_, 1 mM DTT, 5 mM VRC, 1X PMSF, cycloheximide 100 μg/ml, 1% Triton X-100, and 1% sodium deoxycholate) for 30 mins at 4°C followed by centrifugation at 3,000xg for 10 mins at 4°C. Then, the supernatant was then collected and centrifuged at 20,000xg for 10 mins at 4°C. The final supernatant was collected and loaded onto a 30% sucrose cushion in gradient buffer and centrifuged at 31,200 rpm for 1 h at 4°C in a SW61Ti rotor of the Beckman Coulter ultracentrifuge. After centrifugation, the non-polysomal fractions were collected, and the rest of the solution was mixed with dilution buffer by inverting the tubes, followed by centrifugation at 31,200 rpm for 30 min at 4°C in a SW61Ti rotor. The polysome-containing pellet was resuspended in polysome buffer (10 mM Hepes, 25 mM KCL, 5 mM MgCl_2_, 1 mM DTT, 5 mM VRC, and 1× PMSF) and kept for RNA isolation and Western blot(Ghosh *et al*., 2015).

### Immunoprecipitation

Immunoprecipitation was performed as described in earlier papers from our group (Ghoshal *et al*, 2021). Protein G agarose beads (Life Technologies) were used to pulldown exogeneously expressed HA tagged proteins or any other endogenous protein. Beads were washed twice with cold 1X IP buffer (20 mM TRIS–HCl, pH 7.5, 150 mM KCl, 5 mM MgCl_2_, and 1 mM DTT, 1X EDTA free protease inhibitor cocktail) at 2,000xg for 2 mins at 4°C. Beads were then blocked with 5% BSA in lysis buffer for 1 hr at 4°C in a muta-rotator followed by 2X wash with IP buffer. Then, the required amount of antibody was added to the bead, and the tubes were rotated for 4 hrs of incubation at 4°C before lysate addition. The final dilution of antibody was kept 1:100 in every case. Cells were lysed in a lysis buffer (1X IP buffer with 0.5% Triton X-100 and 0.5% sodium deoxycholate) for 30 mins at 4°C. The lysate was not sonicated in these experiments since that could abrogate the fine interactions between the proteins and the miRNAs. Instead, the lysate was subjected to centrifugation at 3000g for 10 mins at 4°C and was added to the bead-antibody mixture and rotated overnight for immunoprecipitation to occur at 4°C. After the reaction, the lysate–bead–antibody mixture was washed with 1X IP buffer thrice, and the final pellet was resuspended in 400 μl IP buffer. The solution was divided into two equal volumes and was kept for protein and RNA estimation. For RNA, Trizol LS was added thrice of the volume, and for protein, 5X SDS dye was added so that the final concentration would be 1X for loading onto a gel.

### Mitochondrial import assay

For the mitochondrial import assay to check the rate of uptake/import of miR-122 within the mitochondria, crude mitochondria were isolated from fed, starved, or treated Huh7 cells according to the protocol already mentioned earlier in the methods section. This mitochondrial population was incubated with synthetic single-stranded miR-122 (10 nM) Eurogentec Inc.) in vitro at 30^0^C for 15 minutes. After the reaction, the residual unimported miRNAs remained in the buffer and were digested with RNase A (5μg/ml) for 15 min at 37°C (Bhattacharyya *et al*, 2003). The mitochondria were then reisolated for estimation and quantification of imported synthetic miR-122 by qRT-PCR.

### TUNEL assay

The TUNEL assay was performed to check the viability of stress-induced cells. For this purpose, the Promega DeadEnd™ Fluorometric TUNEL System kits used, and the assay was done following the manufacturer’s protocol was followed in all cases.

### H_2_O_2_ treatment and autophagy induction

Huh7 cells were cultured on coverslips (for microscopy) or in the multiwell tissue culture plate in a DMEM medium containing 10% FBS. The cells were then treated with 3mM H_2_O_2_ for 10 minutes and revived in media without H_2_O_2_. This was followed by harvesting the cells for microscopy or relevant western blots and RNA isolation. The cells were treated with Lysotracker for 5 minutes for microscopy analysis before fixing them with paraformaldehyde.

### Cell fusion assay

HeLa cells and Huh7 cells were grown separately and transfected with HA-tagged Ago2 and Mito-GFP encoding plasmids along with Cy3-tagged miR-122 (25μm), respectively. The transfected cells were kept for a day after media change and then were co-cultured together on a cover slip. This was followed by the step of fusion of cells mediated by 50% PEG-1500 solution for 10 minutes at 37^0^C followed by wash and revival. For refeeding experiments, refeeding was done for 8 hrs after the fusion step, and then cells were fixed with 4% paraformaldehyde for microscopy.

### Immunofluorescence and imaging

Cells on coverslips were fixed using 4% paraformaldehyde in phosphate-buffered saline (PBS) in the dark at room temperature for 30 min. Coverslips were washed thrice with 1 PBS, blocked, and permeabilized using 1 PBS containing 10% goat serum (Gibco), 1% bovine serum albumin (Affymetrix; USB), and 0.1% Triton X-100 (Calbiochem) for 30 min at room temperature. Primary antibody, diluted in 1 PBS with 1% BSA, was added to cells on coverslips at 4°C overnight in a humid chamber, followed by three washes with 1 PBS for 5 min each time. Alexa Fluor-labeled secondary antibody (1:500; Invitrogen), diluted in 1X PBS with 1% BSA, was added to cells on coverslips. The coverslips were placed in a humid chamber at room temperature for 1 h, and the cells were washed three times with 1X PBS for 5 min each and mounted with Vectashield mounting medium with DAPI (4,6-diamidino-2-phenylindole; Vector).

Cells grown on a six-well tissue culture plate were transfected with 250 ng of GFP–Ago2, GFP-mito, or HA-HuR encoding plasmids(Bhattacharyya *et al*., 2003)for immunofluorescence. The cells were split after 24 hrs of transfection and subjected to various experimental conditions. For immunofluorescence analysis, cells were fixed with 4% paraformaldehyde for 30 minutes in the dark and then washed with PBS. After this, cells were permeabilized and blocked, as mentioned in the previous section. After incubation with primary antibodies in the same buffer at the required dilution overnight at 4°C, subsequent washing was done with PBS three times each for a span of ∼5 mins. Secondary anti-rabbit, anti-rat or anti-mouse-IgG antibodies labeled with Alexa Fluor^®^ 488 dye (green), Alexa Fluor^®^ 594 dye (red), or Alexa Fluor^®^ 647 dye (far red) fluorochromes (Molecular Probes) were used at 1:500 dilutions. After two hours of incubation at 37°C for one hour, followed by washing steps, cells were mounted with Vectashield with DAPI (Vector Labs) and observed under a Zeiss Confocal Imager LSM800.

All standard statistical studies measuring the length, volume, surface area, and ellipticity of mitochondrial networks and individual mitochondria were carried out using the Bitplane Imaris software.

### Statistical analysis

All graphs and statistical analyses were generated in Prism (v5.00) software (GraphPad, San Diego, CA). Nonparametric unpaired and two-tailed paired t-tests were used for analysis keeping a confidence level of 95%. P values of 0.05 or lesser were considered statistically significant, and P values of >0.05 were insignificant. Error bars indicate means standard deviations (SDs)

### *In silico* and bioinformatics analysis

All *in silico* experiments were performed with workstations present at the institute facility. The gene sets that were curated were all through references from earlier reports from various papers as harnessed through text mining. MirTarBase was used for deriving the global miRNA-target gene pairs for both human and mouse studies(Hsu *et al*, 2011) (Huang *et al*, 2020). GSEA genesets were used for the reference pathways and already established organeller gene repertoire for a declared set of proteins from various studies. The Cytoscape version 3.8 was used for various small and large network constructions. Basic parameters like node and edge restrictions were imposed without the involvements of any other add-ons from the software. Further mapping for the network-to-network study was also done in Cytoscape itself. All networks constructed were undirected in nature. Though every edge corresponded to the multiple numbers of evidence for every interaction, we avoided the complications of a weighted network for the sake of simplicity and due to the requirement of only identification of required candidates without any bias.

A cellular scenario for representing various interactions was done with the help of Cell Designer software 4.3, which is available online. No other simulation was done with this software since this study did not investigate cellular flux. A comparative study on the sequences of the miRNAs and detection of consensus sequences alongside a comparison of occupancy was done with the help of MAFFT sequence alignment-based software called Jalview. Other ancillary data plotting was done with the help of online Venn diagram construction software after manually enlisting the data and calculating the percentages and other mathematical derivations. Plots were made with the help of GraphPad Prism 8.

## Data Availability

All data supporting the findings of this study are available from the corresponding authors upon request.

## Acknowledgements

We thank Witold Filipowicz and Gunter Meister for the different constructs used in this study. We thank the Funding body, Dept. of Science and Technology (DST), Govt. of India along with the Council for Scientific and Industrial Research (CSIR), and University Grant Commission (UGC) for the fellowship to SB, SC, and SHC. SNB was supported by The Swarnajayanti Fellowship (DST/SJF/LSA-03/2014-15) from Dept. of Science and Technology, Govt. of India. The work also received support from a High-Risk High Reward Grant (HRR/2016/000093) from Dept. of Science and Technology, Govt. of India, and CEFIPRA project grant 6003-J. SNB is currently supported by the Start-Up Support Grant of the University of Nebraska, USA, and the Lieberman Research Award, Department of Anesthesiology, UNMC support K.M.

## Author Contributions

S.N.B. and K.M. conceived the idea, designed the experiments, and analyzed the data. S.B., S.C., and S.H.C. contributed to the design and planning of the experiments and performed the experiments. K.M., G.H., and S.B. also wrote the manuscript with S.N.B. and analyzed the data.

